# White Matter Microstructural Plasticity Associated with Educational Intervention in Reading Disability

**DOI:** 10.1101/2023.08.31.553629

**Authors:** Steven L. Meisler, John D. E. Gabrieli, Joanna A. Christodoulou

## Abstract

Children’s reading progress typically slows during extended breaks in formal education, such as summer vacations. This stagnation can be especially concerning for children with reading difficulties or disabilities (RD), such as dyslexia, because of the potential to exacerbate the skills gap between them and their peers. Reading interventions can prevent skill loss and even lead to appreciable gains in reading ability during the summer. Longitudinal studies relating intervention response to brain changes can reveal educationally relevant insights into rapid learning-driven brain plasticity. The current work focused on reading outcomes and white matter connections, which enable communication among the brain regions required for proficient reading. We collected reading scores and diffusion-weighted images at the beginning and end of summer for 41 children with RDs who had completed either 1st or 2nd grade. Children were randomly assigned to either receive an intensive reading intervention (*n* = 26; *Seeing Stars* from Lindamood-Bell which emphasizes orthographic fluency) or be deferred to a wait-list group (*n* = 15), enabling us to analyze how white matter properties varied across a wide spectrum of skill development and regression trajectories. On average, the intervention group had larger gains in reading compared to the non-intervention group, who declined in reading scores. Improvements on a proximal measure of orthographic processing (but not other more distal reading measures) were associated with decreases in mean diffusivity within core reading brain circuitry (left arcuate fasciculus and left inferior longitudinal fasciculus) and increases in fractional anisotropy in the left corticospinal tract. Our findings suggest that responses to intensive reading instruction are related predominantly to white matter plasticity in tracts most associated with reading.

## 1. Introduction

Reading disabilities are the most common learning disability (Shaywitz, 1998), impacting as many as 20% of children (Wagner et al., 2020). Formal reading instruction begins at school entry (around 6 years old) for most children in the United States, and readers continue developing their skills in and out of school contexts. However, during extended formal education breaks such as summer vacation, typically a two to three month period in U.S. schools, reading progress typically slows (Cooper et al., 1996; Entwisle et al., 1997; von Hippel et al., 2018). Extended suspension of formal schooling can exacerbate achievement gaps among vulnerable readers who do versus do not participate in reading instruction (Christodoulou et al., 2017) as well as between vulnerable readers and their typically reading peers, as was observed during COVID-19 school disruptions (Kuhfeld et al., 2023). Reading interventions over the summer can halt reading skill loss and even lead to appreciable gains in reading abilities for struggling readers (Christodoulou et al., 2017; Donnelly et al., 2019).

### 1.1 White Matter Supporting Reading

Fluent reading is enabled by coordination of a network of regions in the brain. This system is typically left-lateralized (Murphy et al., 2019), consistent with the frequent left-lateralization of language processing (Enge et al., 2020; Fedorenko et al., 2011; Vigneau et al., 2006). A set of white matter tracts propagate information within this network (Ben-Shachar et al., 2007). Among these tracts are the inferior longitudinal fasciculus (ILF), which connects primary visual cortices to their ipsilateral anterior temporal lobes (Herbet et al., 2018), and the arcuate fasciculus (AF), which connects ipsilateral temporal, parietal, and frontal regions (Catani et al., 2005). Surgical resection and lesion mapping studies of these tracts support their necessity for fluent reading; individuals with damage to these regions exhibit dysfluent reading (Epelbaum et al., 2008; Herbet et al., 2018; Ng et al., 2021; Zemmoura et al., 2015). The ILF is predominantly involved in early processes of word reading, such as orthographic recognition, by carrying primary visual signals to the posterior visual word form area (VWFA; see Dehaene & Cohen, 2011) in the (typically left) ventral occipitotemporal cortex (Bouhali et al., 2014; Lerma-Usabiaga et al., 2018; Yeatman et al., 2013; Yeatman & White, 2021). The ILF’s anterior temporal projections also support semantic processing (Duffau et al., 2013; Herbet et al., 2018; Saur et al., 2008; Shin et al., 2019). The AF’s connections from the anterior VWFA to temporoparietal and frontal regions (e.g., linking Wernicke’s and Broca’s area) are involved in abstracting phonological and higher-order language representations from printed text (Catani et al., 2005; Catani & Mesulam, 2008; Weiner et al., 2017). Other bundles likely support reading as well, including the inferior fronto-occipital fasciculus (ipsilateral frontal-to-occipital connections), superior longitudinal fasciculus (ipsilateral frontal-to-parietal connections), ventral occipital fasciculus (ipsilateral dorsolateral-to-ventrolateral visual cortex connections), and splenium of the corpus callosum (providing interhemispheric visual cortices connections) (Ben-Shachar et al., 2007; Vandermosten et al., 2012; Yeatman et al., 2013), but these bundles have not been as frequent of a focus in reading-related neuroimaging literature compared to the AF and ILF.

### 1.2 Diffusion-Weighted Imaging and White Matter Plasticity

Learning is thought to drive long-term plasticity in white matter (Fields, 2015; Fields et al., 2014; Sampaio-Baptista & Johansen-Berg, 2017; Xin & Chan, 2020). Such changes may manifest as alterations to axonal geometry (e.g., diameter modulations or axonal pruning/branching), myelin remodeling driven by oligodendrocyte proliferation and differentiation, or variations to extra-axonal glial cells and vascular systems (Sampaio-Baptista & Johansen-Berg, 2017). White matter micro- and macro-structural properties can be inferred *in vivo* non-invasively with diffusion-weighted imaging (DWI; Basser et al., 1994). Longitudinal DWI studies have related skill acquisition to white matter changes in behavior-relevant bundles in animals (Blumenfeld-Katzir et al., 2011; Sampaio-Baptista et al., 2013) and humans (Metzler-Baddeley et al., 2017; Scholz et al., 2009), providing evidence for DWI’s utility in quantifying white matter plasticity. Most DWI studies of plasticity have used metrics from the diffusion tensor imaging (DTI) model, including fractional anisotropy (FA) and mean diffusivity (MD). FA measures the degree to which water molecule movement is directionally dependent (varies between 0 - water moves equally as well in all directions, and 1 - water moves only along a single axis), while MD is related to the total magnitude of water movement across all directions. Higher FA and lower MD are thought to indicate well-myelinated white matter (however, see the *Discussion* for limitations surrounding these metrics). Given the high test-retest reliability of DTI measures and modest rates of developmental change (Behler et al., 2021; Wu et al., 2022; Yu et al., 2020), observing significant and rapid changes in these metrics is encouraging in suggesting that white matter alterations are occurring and manifest at a level resolvable by MRI.

### 1.3 Longitudinal Relationships of Reading Performance and White Matter Properties

Reading is an appropriate and educationally relevant domain for investigations of learning-driven neural plasticity. First, reading is a skill that has been socio-culturally introduced too recently to be a product of evolution or natural selection pressure. Second, reading must be explicitly taught, and highly reliable measures exist to gauge reading performance (Torgesen et al., 2012; Woodcock, 2011). Most of the longitudinal neuroimaging literature in reading has been on the order of several months to years. Longitudinal DWI studies of long-term reading development have shown that trajectories of white matter properties and reading skills are significantly linked (Moulton et al., 2019), especially in the left AF, such that increases in tract volume (Myers et al., 2014) and FA (Roy et al., 2024; Van Der Auwera et al., 2021; Wang et al., 2017; Yeatman et al., 2012) accompany improvements in reading among children with diverse reading abilities, although the opposite trend for FA has also been reported for children with reading disabilities (Yeatman et al., 2012). A comparison of illiterate and ex-illiterate adults suggests that developing literacy is associated with higher FA in the left AF (Thiebaut de Schotten et al., 2014). Similarly, higher FA in core reading tracts have predicted subsequently better future reading outcomes in children (Borchers et al., 2019; Davison et al., 2022; Gullick & Booth, 2015), as well as better reading-adjacent skills, such as phonological awareness, among pre-readers (Saygin et al., 2013; Zuk, Yu, et al., 2021). These reports collectively suggest that reading outcomes are related to microstructural changes in white matter that are known to support reading.

In longitudinal studies of reading disabilities, there has not only been a focus on left-hemispheric core reading tracts, which have exhibited lower FA among pre-readers with familial risk of dyslexia (Langer et al., 2017; Vandermosten et al., 2015) and future diagnoses of dyslexia (Vanderauwera et al., 2017), but also on their right hemispheric homotopes. Higher FA in the right superior longitudinal fasciculus has predicted future reading outcomes in children with dyslexia (Hoeft et al., 2011) and longitudinal FA increases in this tract relates to positive reading development in children with familial risk for dyslexia (Wang et al., 2017). These studies suggest the right hemisphere may provide a compensatory mechanism in reading disabilities.

The extant literature implies that white matter infrastructure could have a causal and dynamic relationship with reading outcomes, as opposed to being a static genetically predisposed foundation that reflects individual differences in such outcomes. Indeed, longitudinal designs, as described above, have yielded stronger results than analogous high-powered cross-sectional studies that have suggested little-to-no relationship between DTI measures and individual differences in reading skills (Meisler & Gabrieli, 2022a; Moreau et al., 2018; Roy et al., 2024).

### 1.4 White Matter Plasticity in Reading Intervention

While longitudinal studies of long-term reading development seem to have converged on the importance of left-hemispheric reading circuitry in predicting and tracking reading outcomes, neuroimaging studies focusing on short-term intensive reading instruction (on the order of days-to-weeks) have yielded few and mixed findings on rapid anatomical correlates of reading remediation (as well as functional correlates - (Barquero et al., 2014; Braid & Richlan, 2022; Perdue et al., 2022). Huber et al., 2018 found that decreases in MD across the brain, not limited to core reading circuitry, were related to better reading intervention benefits across participants. One study reported better intervention responses were related to increases in FA only in the anterior left centrum semiovale (Keller & Just, 2009), a broad term for white matter between the corpus callosum and cortical surface which is not considered a part of core reading circuitry. Another study found decreased MD in several left hemispheric regions after reading intervention, although right-hemispheric regions were not reported (Richards et al., 2017). However, a different study concluded that white matter microstructure did not change with reading intervention, but lower right-hemispheric dorsal white matter MD prior to intervention predicted better intervention outcomes (Partanen et al., 2021). In young pre-readers undergoing early literacy training, pre-to-post increases in FA in the left AF and ILF were observed, but these were ultimately attributed to developmental, as opposed to intervention-driven, processes (Economou et al., 2022). In summary, while rapid white matter changes may be observed in a short period of time in the context of intensive reading intervention, it is unclear whether these changes are reproducible, domain-specific (that is, localized to tracts that typically support reading), or dissociable from underlying developmental trajectories. Inconsistent findings could be driven by a variety of factors including publication bias to report positive findings, small sample sizes, and variation in participant characteristics, interventions, and neuroimaging acquisition and analysis protocols (Perdue et al., 2022; Roy et al., 2024; Schilling, Rheault, et al., 2021; Schilling, Tax, et al., 2021; Thornton & Lee, 2000). It is also a possibility that intervention-driven effects are not robust or generalizable, reflecting unique properties of the intervention used or cohort studied.

In the present study, we examined changes in reading skill and white matter microstructure over the course of a six-week summer reading intervention among children with reading disabilities (RD). We focused on properties averaged across all white matter and specifically within 7 white matter tracts: the left AF and ILF as core reading circuitry bundles, their right-sided homotopes as potential compensatory bundles, bilateral corticospinal tracts (CST) as bundles that are not thought to subserve reading, and the splenium of the corpus callosum, which may support reading but has been shown to be microstructurally stable during reading intervention (Huber et al., 2018). We hypothesized observing one of two outcomes: (1) decreases in MD and/or increases in FA in all tracts (besides the splenium) would be related to better intervention responses (e.g., Huber et al., 2018), or (2) this effect would be localized to just the left AF, consistent with multiple studies tracking reading development on longer time scales (Roy et al., 2024; Van Der Auwera et al., 2021; Yeatman et al., 2012).

## 2. Materials and Methods

### 2.1. Ethics Statement

This project was approved by the Massachusetts Institute of Technology’s Committee on the Use of Humans as Experimental Subjects (protocol number: 1201004850). Informed written consent was obtained from parents or legal guardians, while informed written assent was obtained from the participants, who were all minors.

### 2.2. Participants

Participants included in the present study were recruited for a broader overarching study, for which reading (Christodoulou et al., 2017) and gray matter morphometric (Romeo et al., 2018) findings have been previously reported. Forty-one participants passed all inclusion and quality control criteria and were analyzed in the present study (see *Data Inclusion and Quality Control*). All participants were between 7 and 9 years old at the time of enrollment and were entering the summer having completed grades 1 or 2. Inclusion criteria included a history of reading difficulty based on parental report and a manifestation of reading difficulty at study enrollment. In particular, to be included in the study, participants had to have scored “At Risk” or “Some Risk” on the Dynamic Indicators of Basic Early Literacy Skills test (DIBELS; Good et al., 2002) and below the 25th percentile on at least 3 of the 5 following measures: Elision and Nonword Repetition subtests from the Comprehensive Test of Phonological Processing, 2nd Edition (CTOPP-2; Wagner et al., 1999), and the Objects, Letters, and 2-set Letters and Numbers subtests of the Rapid Automatized Naming and Rapid Alternating Stimulus Tests (RAN/RAS; Wolf & Denckla, 2005). Additionally, participants had to score at or above the 16th percentile on the Matrices subtest of the Kaufman Brief Intelligence Test, 2nd Edition (KBIT-2; Kaufman, 2004), which is a measure of nonverbal cognitive ability. All children were native English speakers. Children were recruited from a local partner charter school and the Greater Boston area. Socioeconomic information was collected from parents, who completed the Barratt Simplified Measure of Social Status (Barratt, 2006).

### 2.3. Reading Intervention

Participants were randomly assigned to either receive a reading intervention (*n* = 26) or be placed on a waiting-list (*n* = 15). Comprehensive details of the intervention have been previously described (Christodoulou et al., 2017). Intervention participants completed intensive reading instruction following the *Seeing Stars: Symbol Imagery for Fluency, Orthography, Sight Words, and Spelling* program (Bell, 1997). Instruction was delivered by trained Lindamood-Bell teachers, who rotated classrooms hourly. The program duration was 4 hours per day on 5 days per week for 6 weeks; intervention duration totaled between 100 and 120 hours. Students received small group instruction (3-to-5 students per group) to improve foundational reading skills including phonological and orthographic processing, with an emphasis on visual and orthographic skills. Children recruited from the local partner school received the intervention on-site at their school (*n* = 9), while children recruited from the community-at-large (*n* = 17) received the intervention at a dedicated space at the Massachusetts Institute of Technology.

### 2.4. Outcome Measures

Standardized reading scores were collected from all participants before and after the intervention period, regardless of whether they participated in the intervention. A reading measure proximal to the intervention, the Symbol Imagery Test (SIT), measured orthographic processing in reading (Bell, 2010). During the SIT, participants briefly viewed cards with words or pseudowords for between 2 and 7 seconds and were then asked to report what they were shown. Cronbach’s α values range from 0.86 to 0.88, and the test–retest reliability is 0.95 (Bell, 2010). A relatively distal composite reading index was calculated at each time point by averaging the following four age-standardized reading scores: Sight Word Efficiency (SWE) and Phonemic Decoding Efficiency (PDE) from the Test of Word Reading Efficiency, 2nd Edition (TOWRE-2; Torgeson et al., 1999), and Word Identification (WID) and Word Attack (WA) from the Woodcock Reading Mastery Tests, 3rd Edition (WRMT-3; Woodcock, 2011). Although these measures included orthographic processing, they involved multiple other processes involved in word reading accuracy and fluency. Timed and untimed single word reading skills were measured by SWE and WID, respectively, while timed and untimed pseudoword reading skills were measured by PDE and WA, respectively. For all four subtests, Form A was administered at the beginning of the study, and Form B was administered at the end of the study to avoid practice or familiarity effects. High alternate form reliability has been reported for standardized tests scores on both the WRMT-3 subtests (Word ID: *r* = 0.93, Word Attack: *r* = 0.76; Woodcock, 2011) and the TOWRE-2 subtests (SWE: *r* = 0.90, PDE: *r* = 0.92; Torgesen et al., 2012). Age-normed scores for all tests were defined such that the population mean is 100, with a standard deviation of 15.

### 2.5. Neuroimaging Acquisition

All participants, regardless of intervention site, were scanned at the Athinoula A. Martinos Imaging Center at the Massachusetts Institute of Technology using a 3 Tesla Siemens TimTrio scanner and standard 32 channel head coil. During each session, a T1-weighted (T1w) MPRAGE image was acquired with the following parameters: TR=2.53s, TE=1.64ms, Flip Angle=7°, 1mm isotropic voxels. A diffusion-weighted image (DWI) was acquired with the following parameters: TR=9.3s, TE=84ms, Flip Angle=90°, 2mm isotropic voxels, and 10 b0 volumes followed by 30 non-collinear directions at *b*=700 s/mm^2^. Age-appropriate movies were shown during these scans to increase scan engagement and reduce head motion (Greene et al., 2018). Functional MRI tasks were also collected but are not discussed here. Before the first MRI session, participants were introduced to the MRI by visiting the center’s pediatric mock scanner, which allows children to get acclimated with MRI noise and lying still in the machine, which improves scan compliance (de Bie et al., 2010; Gao et al., 2023).

### 2.6. MRI Preprocessing and Tract Segmentation

MRI preprocessing and tract segmentation were performed according to the longitudinal TRActs Constrained by UnderLying Anatomy (TRACULA) pipeline (Maffei et al., 2021; Yendiki et al., 2011, 2016), as part of *FreeSurfer* version 7.2 (Fischl, 2012; Fischl et al., 2002; Reuter et al., 2012). This method uses longitudinal anatomical priors to produce more plausible tracts compared to creating independent segmentations at each time point (Yendiki et al., 2016), as well as leverages high-quality training data to help inform tract shapes on routine-quality DWI data (Maffei et al., 2021). To achieve this, *FreeSurfer*’s longitudinal processing pipeline (Reuter et al., 2012) was run on each participant’s pre and post T1w images to create an unbiased subject template image (Reuter & Fischl, 2011) using inverse consistent registration (Reuter et al., 2010). Information from this template was used to initialize several steps of the `recon-all` pipeline, such as skull-stripping and anatomical segmentation (Reuter et al., 2012).

DWI volumes from each image were aligned to the first b0 image in that scan. The *b*-matrix was rotated accordingly (Leemans & Jones, 2009). DWI images were corrected for motion and eddy currents with FSL’s `eddy` command (Andersson & Sotiropoulos, 2016). Information from this process was used to generate four measures of head motion and image quality that inform the “total motion index” (Yendiki et al., 2014): mean volume-by-volume head rotation, mean volume-by-volume head translation, proportion of slices with signal dropout, and severity of signal dropout. The diffusion tensor was fitted using FSL’s `dtifit`. Mean diffusivity (MD) and fractional anisotropy (FA) were derived from the tensor. A GPU-accelerated ball-and-stick model was fit for each DWI image (Behrens et al., 2007; Hernández et al., 2013; Jbabdi et al., 2012). At each time point, a registration was computed between the diffusion-weighted image and T1w image (native space) using an affine boundary-based registration algorithm (Greve & Fischl, 2009). This transformation was used to bring anatomical segmentations into DWI space. The DWI-to-T1w and T1w-to-template registrations were multiplied to get a DWI-to-template transformation. Information from high-resolution 7T training data (Maffei et al., 2021) was used to estimate endpoint ROIs and pathways for white matter tracts in each participant’s template space images. These data were then brought back into the native DWI space of each time point. The DWI ball-and-stick model and tract anatomical priors were used to calculate the probability density of each pathway. From these, we collected the average MD and FA from the cores of our tracts of interest (`[MD|FA]_Avg_Center`) to mitigate concerns of noise and partial volume effects from fibers branching towards the exterior and extremities of the bundles. These tracts included the bilateral AF, ILF, and CST, as well as the splenium of the corpus callosum (*Figure 1*). Additionally, at each time point, we calculated the laterality index of microstructural measures among bilateral tracts ([L-R]/[L+R]), as well as the average of the microstructural measures within the *FreeSurfer*-produced white matter segmentation mask. This average, unlike a whole-brain average, does not covary with the proportions of gray-to-white matter.

**Figure 1:**
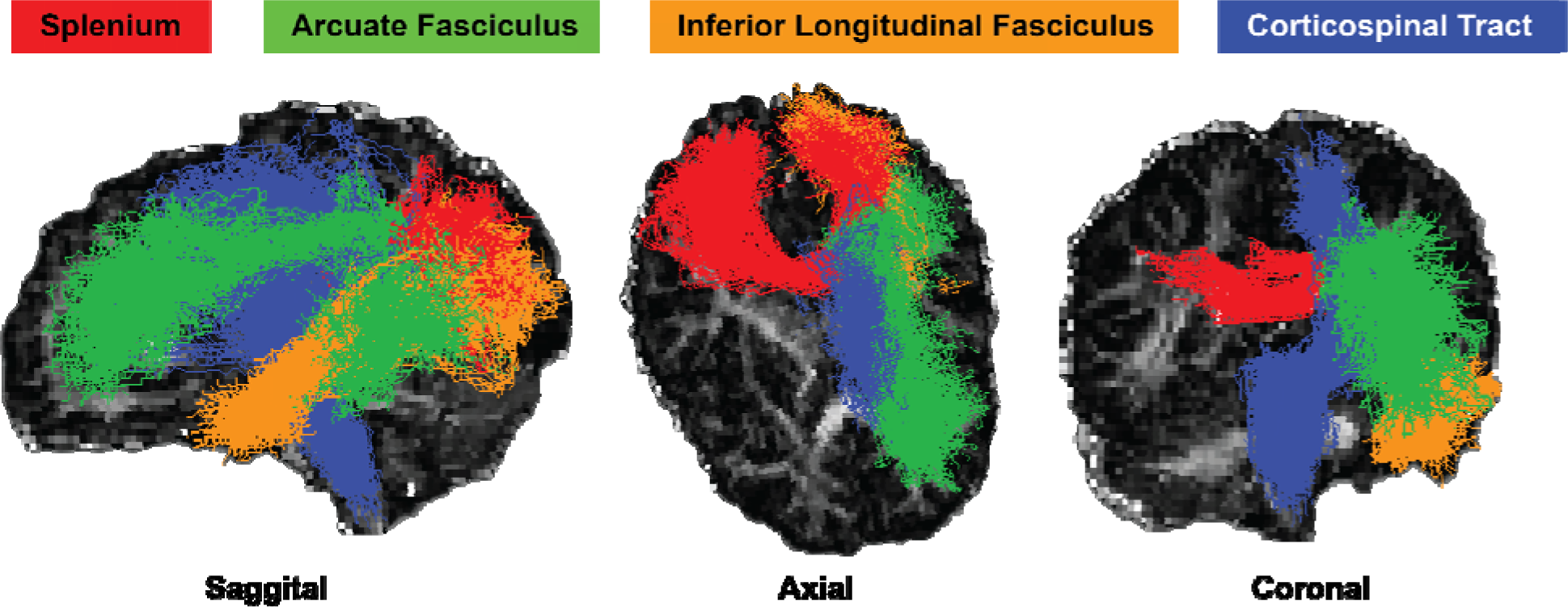
Tracts produced by TRACULA analyzed in the present study, overlaid on top of a fractional anisotropy image. Only the left hemispheric bundles are visualized for bilateral tracts. Pictured data come from a single representative participant.

### 2.7. Statistics and Analysis

Analyses were prepared, run, and visualized using Python packages Pandas 1.3.2 (McKinney, 2011), Statsmodels 0.13.5 (Seabold & Perktold, 2010), and Seaborn 0.12.1 (Waskom, 2021), respectively. We made a dataframe that contained the following phenotypic fields for each subject: ages at each scan (in months), sex (binary categorical factor), and reading measures at pre and post time points (as well as their longitudinal pre-to-post differences). For each tract, the pre and post MD and FA metrics were added to the dataframe, along with their longitudinal pre-to-post differences. Laterality indexes of microstructural measures and their longitudinal differences for bilateral tracts were also added. Microstructural measures averaged across white matter and their longitudinal differences were additionally added. The total motion indices, calculated separately for each time point, were also added to the dataframe (see *Data Inclusion and Quality Control*).

We used ordinary least squares linear models to run multiple regressions, allowing us to relate reading measures to white matter microstructure while controlling for confounds. For the primary analyses, we created models to relate pre-to-post changes in a given tract microstructural measure (dependent variable) to the longitudinal difference in a reading measure across all participants (independent variable), with nuisance regressors for sex, age at first scan, and motion indices at both time points. We also ran a series of related supplemental analyses to provide additional context for the analysis. These included: (1) models with radial and axial diffusivities (RD and AD, respectively) as the primary DWI metric (*Figure S1*); (2) models using just participants who completed the reading intervention (*Figure S3*); and (3) cross-sectional models at each time point (with age and motion confounds being derived from the particular time point) (*Figure S4*). For each model, the effect size (ΔR^2^_adj_) was calculated as the difference in adjusted R^2^ coefficients between that from the full model and from a reduced model without the reading score predictor of interest. A family of tests was considered as the set of tests across tracts for a given reading measure and microstructural metric. This included up to 11 tests (3 bilateral tracts with their laterality indexes, the splenium, and the average white matter). Benjamini-Hochberg false-discovery rate (FDR) correction for multiple hypotheses (Benjamini & Hochberg, 1995) was performed within each family of tests.

### 2.8. Data Inclusion and Quality Control

A total of 153 children were recruited as part of a broader study. Fifty-two participants had anatomical and DWI scans at both time points and were able to complete the neuroimaging processing pipeline without errors. Forty-four of the remaining participants had the necessary phenotypic data. As a quality assurance metric, we computed the total motion index (TMI; Yendiki et al., 2014). TMI is related to four measures: rotation, translation, signal dropout prevalence, and signal dropout severity. For each scan, we calculated each motion metric’s difference from the study population mean for the given time point, divided by the interquartile range of the metric. The TMI for each scan is the cumulative sum of these calculations across the four motion metrics. Three subjects had outliers in TMI at either time point and were excluded. For further quality assurance, we confirmed that no remaining participant had any tract-averaged FA lower than 0.3, which could indicate some combination of white matter disorganization and partial volume effects from a tract branching into significant amounts of gray matter or CSF. Thus, a total of 41 subjects (26 who received an intervention, and 15 in the non-intervention group) were analyzed in the present study.

We conducted a power analysis, in which we used an estimated effect size of |*r*| = 0.40. This corresponds to the relationship between mean diffusivity and changes in TOWRE reading scores observed in the left AF and ILF in a related study (see *Table 1* in Huber et al., 2018 for reference). To resolve that effect at α = 0.05 and power of 0.8, one would need *n* = 34 participants, as calculated by the *G*Power* software version 3.1 (Faul et al., 2009), which our sample exceeds.

**Table 1:** Cognitive and phenotypic summary statistics Values are provided as Mean (SEM). All cognitive measures are age-standardized. Two-sample t-tests were used to test for differences between groups, with the exception of χ^2^-tests for sex and handedness distributions. * denotes p < 0.05. Abbreviations: SES - Socioeconomic Status; KBIT - Kaufman Brief Test of Intelligence; SIT - Symbol Imagery Test; TOWRE - Test of Word Reading Efficiency; SWE - Sight Word Efficiency; PDE - Phonemic Decoding Efficiency; WJ - Woodcock Johnson.

| Participants | All<br>(n = 41) | Intervention<br>(n = 26) | Non-Intervention<br>(n = 15) | Effect Size |
| --- | --- | --- | --- | --- |
| Age at first scan (months) | 94.85 (1.16) | 95.00 (1.32) | 94.60 (2.29) | $d = 0.053$ |
| Sex (M/F) | 26 / 15 | 16 / 10 | 10 / 5 | $\Phi = 0.051$ |
| Handedness (L/R) | 7 / 34 | 7 / 19* | 0 / 15* | $\Phi = 0.345^*$ |
| SES (years of parental education) | 17.89 (0.41) | 18.00 (0.50) | 17.70 (0.75) | $d = 0.112$ |
| KBIT Matrices | 101.7 (2.03) | 100.3 (2.41) | 104.2 (3.70) | $d = 0.301$ |
| SIT (Pre/Post/Diff) | 89.76 (1.77) /<br>93.20 (1.69) /<br>3.439 (2.06) | 88.62 (2.15) /<br>97.07 (1.87)* /<br>8.462 (2.15)* | 91.73 (3.11) /<br>86.47 (2.53)* /<br>-5.267 (3.21)* | $d = 0.275$ /<br>$d = 1.102^*$ /<br>$d = 1.191^*$ |
| Composite Reading Index (Pre/Post/Diff) | 83.21 (1.28) /<br>81.11 (1.34) /<br>-2.101 (0.957) | 82.37 (1.51) /<br>82.34 (1.57) /<br>-0.029 (1.27)* | 84.67 (2.34) /<br>78.97 (2.42) /<br>-5.694 (0.859)* | $d = 0.280$ /<br>$d = 0.395$ /<br>$d = 1.102^*$ |
| TOWRE SWE (Pre/Post) | 83.49 (1.78) /<br>80.46 (1.99) | 83.46 (2.23) /<br>80.81 (2.61) | 83.53 (3.06) /<br>79.87 (3.11) | $d = 0.006$ /<br>$d = 0.073$ |
| TOWRE PDE (Pre/Post) | 79.88 (1.40) /<br>77.63 (1.51) | 78.46 (1.64) /<br>79.42 (1.74) | 82.33 (2.48) /<br>74.29 (2.70) | $d = 0.439$ /<br>$d = 0.552$ |
| WJ Word ID (Pre/Post) | 83.68 (1.45) /<br>82.68 (1.46) | 83.50 (1.79) /<br>83.69 (1.89) | 84.00 (2.57) /<br>80.93 (2.27) | $d = 0.053$ /<br>$d = 0.295$ |
| WJ Word Attack (Pre/Post) | 86.00 (1.70) /<br>83.80 (1.43) | 84.32 (2.06) /<br>85.42 (1.52) | 88.80 (2.89) /<br>81.00 (2.84) | $d = 0.421$ /<br>$d = 0.489$ |

## 3. Results

### 3.1. Cognitive and Phenotypic Data

Phenotypic summary statistics for the participant cohort are provided in *Table 1*. Of note, the intervention and non-intervention groups were matched in sex (χ^2^ test, *p* > 0.7), but not handedness (χ^2^test, *p* < 0.05). We did not include handedness as a regressor in our models due to a lack of evidence of handedness-related asymmetry in white matter microstructure (Jang et al., 2017; López-Vicente et al., 2021). The two groups were also matched in age, non-verbal intelligence (KBIT), socioeconomic status, and reading performance across all subtests cross-sectionally at each time point (two-sample *t*-test, *p* > 0.1 across all tests), with the exception of the intervention group having a significantly higher SIT score post-intervention (two-sample *t*-test, *p* < 0.005). Demonstrating the efficacy of the reading intervention, the intervention group showed larger longitudinal pre-to-post differences in the SIT and composite reading index (two-sample *t*-test, *p* < 0.002 across both tests), driven by the non-intervention group regressing in both measures and the intervention group improving on the SIT and maintaining scores on the composite reading index (*Figure 2*). These results are consistent with what was observed in the larger cohort from which the present subset was derived (Christodoulou et al., 2017).

**Figure 2:**
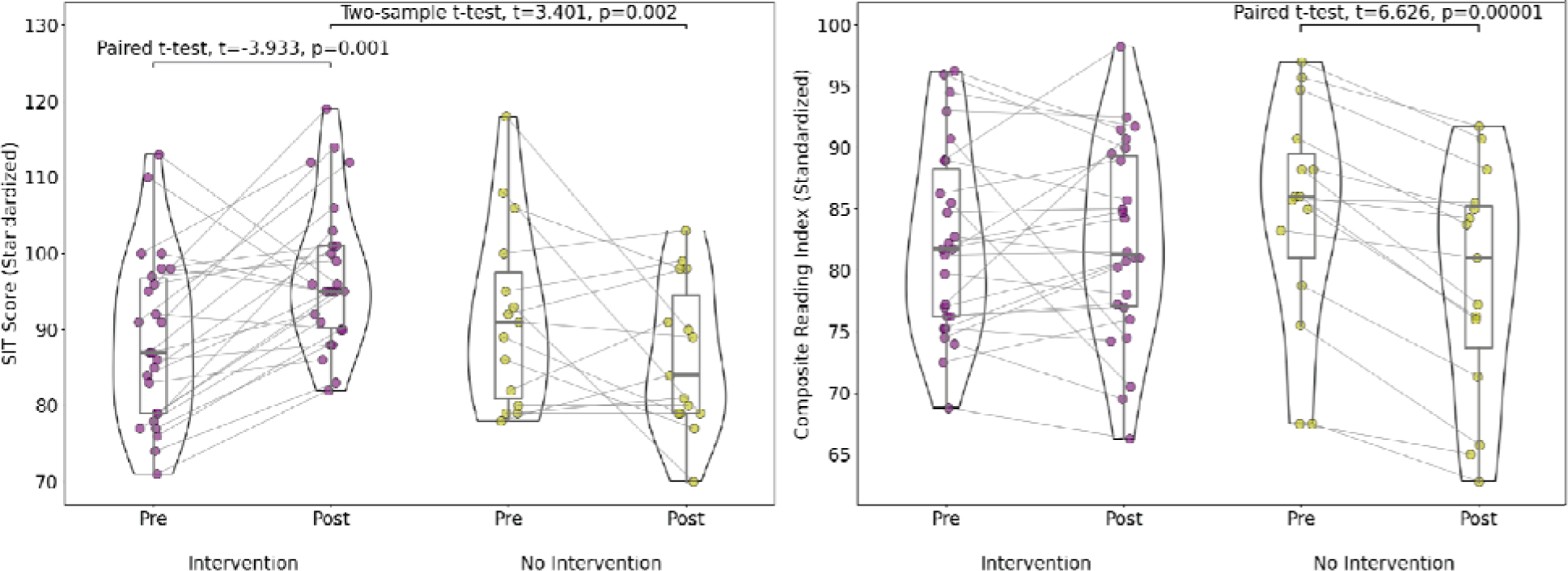
Changes in Symbol Imagery Test (SIT; left) and Composite Reading Index (right) scores for intervention (purple) and non-intervention (yellow) participants. Paired t-tests were used to compare pre and post scores within groups, and two-sample t-tests were used to compare scores at a given time point across groups. Significant tests (p < 0.05) are annotated in the figure.

### 3.2. Relationship Between Changes in White Matter Microstructure and Reading Scores

Across all participants, pre-to-post decreases of MD in the left AF and left ILF were related to improvements in SIT scores over the summer (*Table 2, Figure 3*). SIT score trajectories accounted for ∼9% of variance among MD changes in the left AF, and ∼16% of MD difference variance in the left ILF. Additionally, longitudinal differences in SIT scores accounted for ∼21% of variance in changes in ILF MD laterality, following a similar trend of decreasing leftward laterality of MD relating to improvements in reading. However, similar effects were not present when considering the composite reading index. Similar patterns to MD results were also observed when using radial diffusivity as the DWI metric of interest (*Figure S1*). Decreasing splenium MD was marginally correlated with improvements in both reading measures (*p* < 0.1), with each test score accounting for ∼5% of variance in microstructure. Increasing FA in the left CST (*p* < 0.05) and to a lesser extent the left AF (*p* < 0.1) were related to improvements in SIT scores, but not the composite reading index (*Figure 3, Table 3*). Changes in white matter average FA and MD did not relate to changes in either reading measure (and these metrics were not different between groups before or after the intervention, *Figure S2*). After multiple comparison correction, the models relating changes in SIT scores to differences in the ILF MD laterality (*p_FDR_* = 0.037) and left ILF MD (*p_FDR_* = 0.056) remained statistically significant or marginally significant.

**Figure 3:**
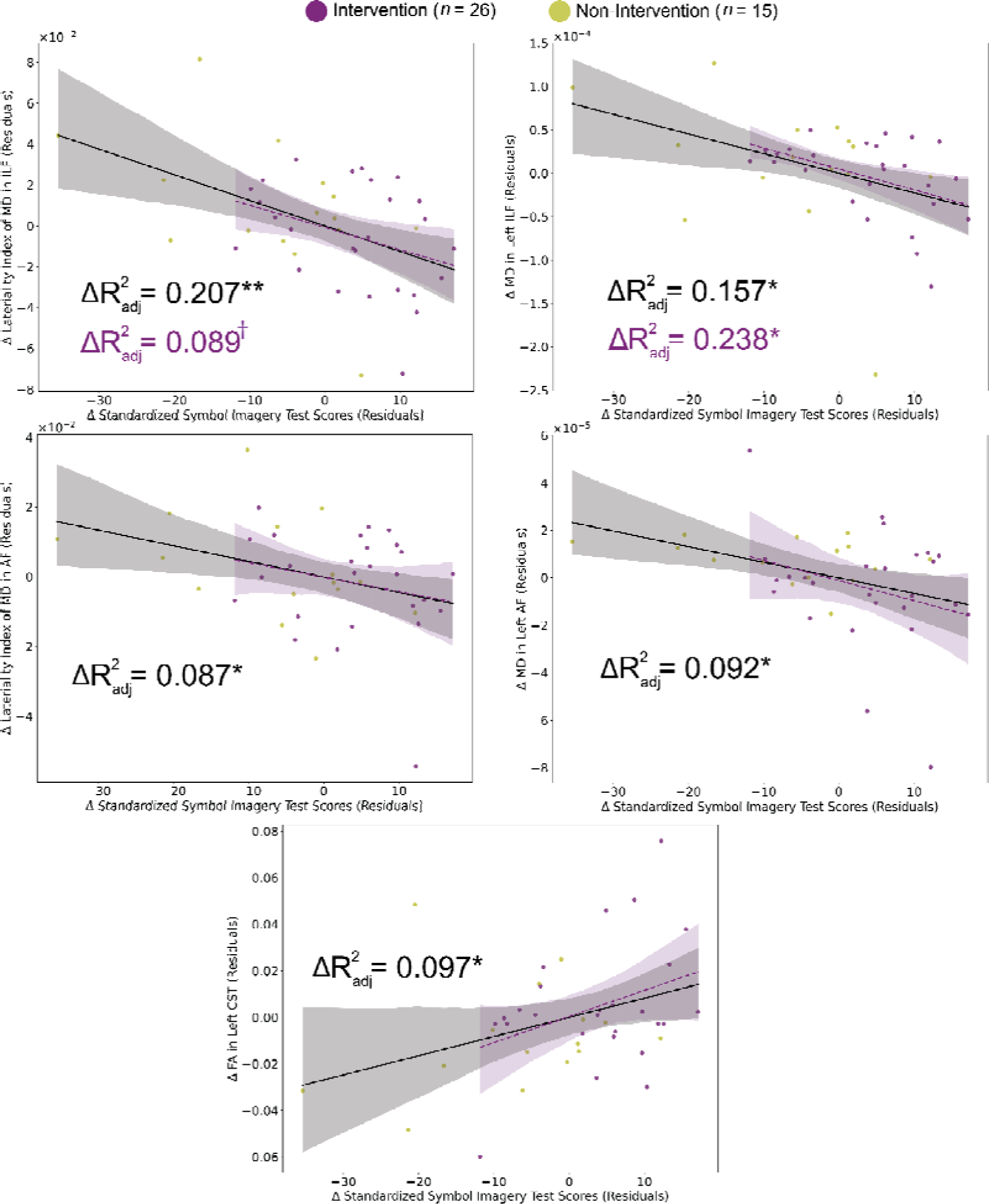
Partial regression plots relating changes in tract microstructure to changes in standardized SIT scores. Confounds included age at first scan, sex, and motion indices at each time point. Values on axes are residuals after accounting for nuisance regressors in the model. Models with an uncorrected p < 0.05 across all participants are shown. For MD, these include models of the left ILF and ILF laterality index (top), and left AF and AF laterality index (middle). For FA, this includes the left CST (bottom). No test reached this threshold with the composite reading index. Purple dots represent intervention participants, and yellow does represent non-intervention participants. The black solid lines and effect sizes represent the fit across all participants, and the purple dashed lines and effect sizes represent the best when considering only intervention participants. †: p < 0.1, *: p < 0.05, **: pFDR < 0.05. Abbreviations: AF - Arcuate Fasciculus; ILF - Inferior Longitudinal Fasciculus; CST - Corticospinal Tract.

**Table 2:** Multiple regression outcomes relating changes in age-standardized reading scores and tract MD across all participants. * denotes p < 0.05, † denotes p < 0.1. Abbreviations: AF - Arcuate Fasciculus; ILF - Inferior Longitudinal Fasciculus; CST - Corticospinal Tract; SIT - Symbol Imagery Test.

| Tract | SIT Score |  |  |  | Composite Reading Index |  |  |  |
| --- | --- | --- | --- | --- | --- | --- | --- | --- |
| | $\beta$ [95% CI] | $\Delta R^2_{adj}$ | $p$-value | $p_{FDR}$ | $\beta$ [95% CI] | $\Delta R^2_{adj}$ | $p$-value | $p_{FDR}$ |
| **White Matter Average MD** | -0.198<br>[-0.559, 0.163] | 0.006 | 0.274 | 0.381 | -0.022<br>[-0.338, 0.294] | -0.025 | 0.889 | 0.889 |
| **Left AF** | -0.385<br>[-0.746, -0.024] | 0.092 | 0.037* | 0.130 | -0.219<br>[-0.541, 0.104] | 0.024 | 0.177 | 0.650 |
| **Right AF** | 0.049<br>[-0.334, 0.431] | -0.026 | 0.798 | 0.810 | 0.080<br>[-0.249, 0.408] | -0.021 | 0.625 | 0.707 |
| Laterality Index AF | -0.370<br>[-0.735, -0.005] | 0.083 | 0.047* | 0.130 | -0.260<br>[-0.581, 0.060] | 0.046 | 0.108 | 0.596 |
| **Left ILF** | -0.480<br>[-0.839, -0.121] | 0.157 | 0.010* | 0.056† | -0.079<br>[-0.417, 0.260] | -0.023 | 0.641 | 0.707 |
| **Right ILF** | 0.050<br>[-0.330, 0.430] | -0.026 | 0.792 | 0.810 | 0.104<br>[-0.222, 0.430] | -0.016 | 0.520 | 0.707 |
| Laterality Index ILF | -0.540<br>[-0.889, -0.193] | 0.207 | 0.003* | 0.037* | -0.186<br>[-0.520, 0.148] | 0.008 | 0.267 | 0.655 |
| **Left CST** | -0.197<br>[-0.558, 0.165] | 0.005 | 0.277 | 0.381 | -0.073<br>[-0.388, 0.243] | -0.201 | 0.643 | 0.707 |
| **Right CST** | -0.193<br>[-0.539, 0.154] | 0.006 | 0.267 | 0.381 | -0.156<br>[-0.454, 0.143] | 0.002 | 0.298 | 0.655 |
| Laterality Index CST | 0.046<br>[-0.338, 0.430] | -0.027 | 0.810 | 0.810 | 0.144<br>[-0.183, 0.471] | -0.006 | 0.378 | 0.692 |
| **Splenium** | -0.292<br>[-0.624, 0.039] | 0.046 | 0.082† | 0.180 | -0.265<br>[-0.549, 0.019] | 0.054 | 0.066† | 0.596 |

**Table 3:**
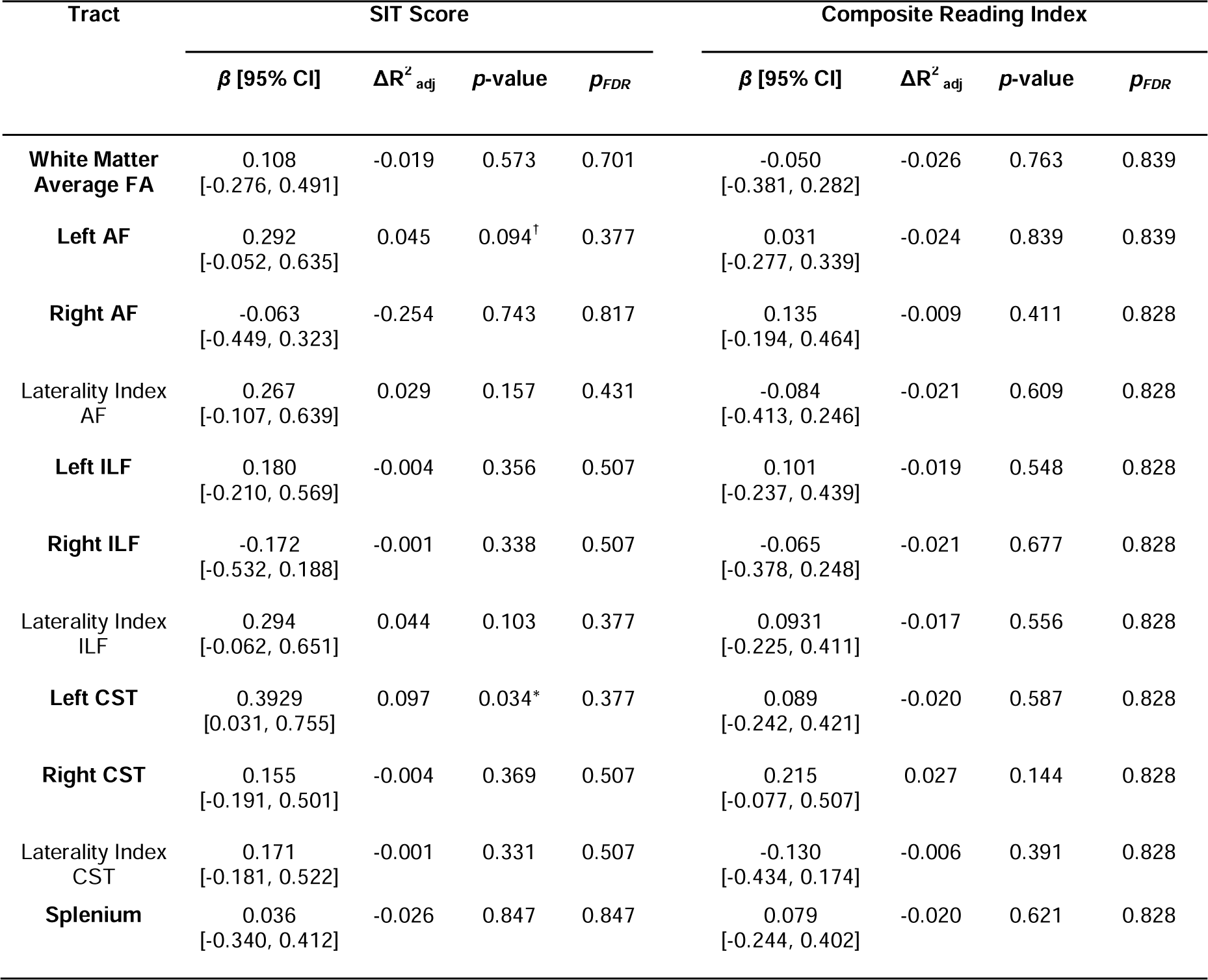
Multiple regression outcomes relating changes in age-standardized reading scores and tract FA across all participants. * denotes p < 0.05, † denotes p < 0.1. Abbreviations: AF - Arcuate Fasciculus; ILF - Inferior Longitudinal Fasciculus; CST - Corticospinal Tract; SIT - Symbol Imagery Test.

When running the same models on only the 26 participants who completed the intervention (*Figure S3*), significant relationships remained between pre-to-post decreases in MD in the left ILF and improvement in SIT scores (*Figure 3*, *p* < 0.05, ΔR^2^_adj_ = 0.238) and between pre-to-post decreases in MD in the splenium and improvement in composite reading index scores (*p* < 0.1, ΔR^2^_adj_ = 0.093). Additionally, improvements in SIT scores were associated with decreases in white matter average MD (*p* < 0.05, ΔR^2^_adj_ = 0.113) and with decreasing FA in the right ILF (*p* < 0.1, ΔR^2^_adj_ = 0.123). The relationship between MD decreases in the left ILF and improvements in SIT scores remained marginally significant (*p* < 0.1; ΔR^2^_adj_ = 0.098) after including white matter MD as an additional covariate *post-hoc*. None of these supplementary models remained significant at α = 0.05 after FDR correction for multiple hypotheses.

## 4. Discussion

In the present study, we investigated whether changes in white matter microstructure were related to changes in reading skill during the summer among 41 children with reading disabilities. Reading ability trajectories varied on a wide spectrum, including score regression (the “summer slump”) and intervention-driven improvement. We focused on 7 tracts within and outside of core reading circuitry, using two microstructural measures (FA and MD), and two reading measures. One reading measure, the SIT, was closely related to the orthographic focus of the intervention, while a separate composite reading index was more distal to the intervention and indexed a broader range of reading-related skills (e.g., phonological awareness and rapid automatized naming). We found that longitudinal decreases in MD (and leftward laterality of MD) were related to improved SIT scores in left-hemisphere core reading circuitry (the AF and ILF). Longitudinal increases in FA in the left CST were also related to improved SIT scores. Notably, none of these associations were present when considering the composite reading index, and only the relationship between improving SIT scores and decreasing leftward laterality of MD in the ILF was significant after FDR correction.

We originally hypothesized that we would see a relationship between improving white matter microstructure (lower MD and/or higher FA) and improving reading scores in either just the left AF or more globally. While neither of these hypotheses were supported, the pattern of results suggests that intervention effects were strongest (and in the hypothesized direction) within core reading circuitry when considering the reading measure most related to the intervention. The specificity of effects to core reading circuitry is supported by the use of laterality index (which tends to be unrelated to global trends) and the mostly null results in models using white matter averaged microstructural measures. That ILF plasticity was most related to reading trajectories, compared to changes in the AF, might reflect the higher relative emphasis on orthographical and visual training in the intervention program (as opposed to processing phonological representations of print, which would be subserved by the AF). This might also explain changes observed in the splenium, which is thought to subserve more basic visual processes. The nature of the intervention may influence what brain structures are affected; for example, phonological-based instruction may have effects localized to brain structures supporting phonological processing (Eden et al., 2004). Similarly, this pattern of results we observed might be representative of the relatively high reliance on visual and orthographic processing in early readers still learning the basics of decoding for reading (Badian, 2001).

The design of our study most closely resembles that of Huber et al., 2018, but with some important differences. The same intervention curriculum (*Seeing Stars*) was used in both studies, with similar instruction hours per week, but the present study had a shorter 6-week intervention period compared to the 8 weeks in Huber et al., 2018. Additionally, students in the present study were taught in small group settings, while a 1-on-1 approach was used in Huber et al., 2018. The longer intervention duration and more intense individualized instruction may have been factors that led to children in that study improving on their composite reading measure (composed of the same reading tests as in the present study), as opposed to only maintaining scores as found in the present study. Both studies had similar cohort sizes (41 and 43 children). However, the present study had participants within a narrower age-range of 7-9 years compared to 7-12 years old in Huber et al., 2018. Additionally, all participants in the present study were diagnosed with a reading disability, while 10 children in Huber et al., 2018 were typical readers. Both studies found that changes in MD in the left AF and ILF were related to reading outcomes over the course of the summer. However, Huber et al., 2018 found more widespread plasticity that did not include the splenium, while our cohort exhibited more domain-specific plasticity that included the splenium (*p* < 0.1). Beyond the contrasts between studies explained above, additional variation in data acquisition, processing, and statistical techniques likely contributed to differences in results between studies (Schilling, Rheault, et al., 2021; Schilling, Tax, et al., 2021).

It is encouraging that both the present study and Huber et al., 2018 found intervention effects in the left AF and ILF, which provides converging lines of evidence suggesting that intense educational instruction influences reading-relevant white matter tracts, albeit with different findings about plasticity occurring in a broader range of tracts. Future studies ought to address similar questions in different contexts to evaluate the reproducibility and generalizability of these findings. Presently, there are not enough extant studies on longitudinal neuroanatomical correlates of reading intervention (in either gray or white matter) to perform meaningful meta-analyses. In functional MRI, a meta-analysis of 8 studies with longitudinal neuroimaging and cognitive scores concluded that there were no consistent locations where longitudinal changes in reading-invoked BOLD signal and intervention response covaried (Perdue et al., 2022). However, individual studies, including those that may have only contained one session of neuroimaging either prior or after intervention, have found es of intervention response both in putative reading regions as well as more globally (reviewed in Barquero et al., 2014; Braid & Richlan, 2022; Perdue et al., 2022). One of the few studies of gray matter morphometric correlates of intervention response was conducted on the same participant pool as in the present study (Romeo et al., 2018). This study concluded that children who improved exhibited significant cortical thickening in brain regions including the left middle temporal gyrus, right superior temporal gyrus, and bilateral middle-inferior temporal cortex, inferior parietal lobule, precentral cortex, and posterior cingulate cortex. While some of these regions comprise left-lateralized core reading areas, others extend globally beyond the reading network. Considering these spatially distinct patterns of results for gray and white matter, it is unclear yet how to coincide different anatomical and functional measures of plasticity in response to reading intervention.

In exploratory analyses, we created cross-sectional models to investigate whether white matter microstructure and reading skills were associated at each time point (*Figure S4*). Lower MD and higher FA in the left ILF were associated with better reading scores at the beginning of the summer, consistent with the left ILF’s critical role in supporting reading. We also found that the right ILF and right CST microstructure had significant associations with reading scores. Notably, this is not consistent with a previous report showing an inverse relationship between right ILF FA and reading outcomes among children with reading disabilities (Banfi et al., 2018), and in the present study, longitudinal microstructural trajectories in these right-sided homotopes were not linked with reading score changes. This might suggest that right-lateralized white matter serves as a static compensatory agent that reflects early reading outcomes in reading disabilities (e.g., Zuk, Dunstan, et al., 2021), but does not dynamically change with reading instruction. However, given the small sample size and limited power of cross-sectional designs, this should be interpreted with caution.

The significance of the models including the CST, both cross-sectionally and longitudinally, were unexpected given its seeming lack of a role in reading as a primary motor tract. Although effect sizes for these models were appreciably lower than those for the left ILF and AF, this suggests that intervention effects could still be detected to some extent outside of reading circuitry. The CST is not often focused in studies of reading abilities, but one study found that volumes of bilateral CST were informative in predicting dyslexia diagnoses in children (Cui et al., 2016). Additionally, other studies have found that FA in the left CST predicted future phonological skills (a critical pre-reading ability) in kindergarten (Zuk, Yu, et al., 2021), correlated with phonological processing in preschoolers (Walton et al., 2018), and corresponded with phonological encoding abilities in adults with brain damage (Han et al., 2016). This suggests that the left CST could be co-opted into reading circuitry in populations with deficient or not-fully developed language abilities, albeit its role in this context is not clear. This deviates from the more frequent focus of compensation from right-sided reading circuitry homotopes such as the right ILF and AF. However, speech and reading difficulties tend to co-occur (Catts, 1993; Hayiou et al., 2010), and the left CST may be important for speech, evidenced by microstructural deficiencies in pre-term children with poor oromotor outcomes (Northam et al., 2012) and stuttering populations (Connally et al., 2014; Kronfeld-Duenias et al., 2016). It is also a possibility that the CST reconstructions largely intersected with the nearby corticobulbar projections, as tractography is prone to overlaps (Schilling et al., 2022). Corticobulbar projections innervate cranial nerves that support head and neck muscles, and properties of corticobulbar tracts have also been related to speech and language outcomes in preterm adolescents (Northam et al., 2019).

Associations between white matter microstructure and reading outcomes were found in relation to a proximal measure of orthographic skill, but not distal measures that included multiple processes related to reading words and pseudowords. The proximal outcome measure (SIT) involved rapid identification of letter sets and was thus directly aligned with the *Seeing Stars* intervention program’s emphasis on orthographic skills. We also expected but did not find statistically significant associations between white matter microstructure and the distal reading composite that included four single word reading measures (timed and untimed, real and pseudo-words). Previous research on this intervention reported the largest effect sizes (Cohen’s *d*) for the SIT measure (1.32), but more modest effect sizes for distal single word reading measures that constituted the reading composite score (Untimed word reading (WRMT WID): 0.96; untimed pseudoword reading (WRMT WA): 0.87; timed word reading (TOWRE SWE): 0.19; timed pseudoword reading (TOWRE PDE): 1.08; Christodoulou et al., 2017). The association between plasticity in white matter microstructure and change in SIT scores may reflect this specific substantial change, perhaps reflecting a minimal threshold of intervention impact that ties to structural plasticity.

The pattern of results in the present study favoring stronger effects in MD than FA imply that the microstructural changes accompanying reading development are related to extra-axonal factors, such as neurite density and CSF volume (Beaulieu, 2002; Genc et al., 2017), as opposed to axonal factors such as myelination, orientation coherence, and axonal density (Friedrich et al., 2020). This is consistent with other DWI studies of reading intervention (Huber et al., 2018, 2021). Our supplemental analyses, which showed that our results with MD were similar to models with radial (but not axial) diffusivity (*Figure S1*), support the idea that improvements in reading might be limited to factors that restrict water movement in extracellular space, such as increased density of axons (Winklewski et al., 2018). However, multimodal research at various spatial and temporal resolutions will need to be reconciled to perform the nontrivial task of ascribing such changes to biophysical mechanisms (Jelescu et al., 2020). While higher FA and lower MD are often thought to reflect more “healthy” white matter, these metrics are biologically unspecific due to the variety of factors that can influence diffusion of water in voxel-sized regions and the confounding impact of crossing fibers (De Santis et al., 2014; Jones et al., 2013), which can impact as many as 90% of white matter voxels (Behrens et al., 2007; Jeurissen et al., 2013). FIber-specific measures such as quantitative anisotropy (Yeh et al., 2013) and fixel-based metrics (Raffelt et al., 2017), and multicompartmental models such as NODDI (Zhang et al., 2012), can provide higher biological specificity and have shown promise in better resolving brain-behavior relationships in studies of reading abilities (Koirala et al., 2021; Meisler & Gabrieli, 2022b; Sihvonen et al., 2021). Unfortunately, the low angular resolution and weak single-shelled diffusion weighting of the present DWI acquisition scheme were not well-suited for these more novel approaches (Genc et al., 2020), effectively limiting us to using DTI metrics. Outside of DWI, related white matter neuroimaging sequences, such as myelin water imaging and quantitative T1 imaging, may provide more targeted insights into learning-driven plasticity in reading (Economou et al., 2023; Huber et al., 2021). Future longitudinal studies should consider incorporating these techniques.

Our results should be considered in the context of additional limitations. First, the only test that survived multiple comparison correction was the one that suggested decreasing leftward laterality of MD in the ILF was related to improving SIT scores (and the analogous model concerning only the left ILF was marginally significant at *p_FDR_*= 0.056). Despite this, we believe the strong effect sizes achieved by many of the models, even those that did not survive FDR correction, are noteworthy given the typically small effect sizes observed in analogous cross-sectional analyses (e.g., ∼3% variance explained, as observed in Meisler & Gabrieli, 2022b). The small amount of within-participant data (two time points) precluded us from running more statistically sophisticated models, such as linear mixed-effect models as in (Huber et al., 2018). This also limited our ability to infer when microstructural changes occurred over the course of the intervention and characterize the temporal relationship between changes in reading and tract properties (in other words, whether tract microstructural changes preceded changes in reading scores, or vice-versa). To address these points, future longitudinal studies of reading intervention should strive to contain at least 3 sessions of data collection (King et al., 2018). Our relatively small sample size of 41 reflects the challenges of collecting acceptable quality cognitive and multi-modal MRI data in children with learning disabilities undergoing intervention, which are particularly amplified for longitudinal studies (Davis et al., 2022). Even with these limitations, we found that white matter microstructural plasticity, predominantly in core reading circuitry, was related to changes in reading abilities over the summer in the context of short-term intensive educational intervention.

## Data and Code Availability

Due to language used in the consenting process, we are not permitted to publicly share subject MRI images. Images may be privately distributed upon reasonable request. We share a CSV containing all necessary data to replicate the present results, as well as the code to recreate the analyses and figures. All instructions and code for processing data and running the statistical analyses can be found at https://github.com/smeisler/Meisler_ReadingInt_DWI. To execute the *FreeSurfer* workflows, we ran a Docker container containing *FreeSurfer* 7.2 and *FSL* 6.0.4 with Singularity (3.9.5) (Kurtzer et al., 2017). The container can be collected with either `docker pull amirro/tracula:latest` or `singularity build tracula_container.img docker://amirro/tracula:latest`. Development of these software may introduce improvements and bug fixes that should be used in future research, so we encourage using the latest stable releases.

## Author Contributions

**Steven L. Meisler:** Conceptualization, Formal analysis, Funding acquisition, Methodology, Software, Visualization, Writing - original draft, Writing - review & editing. **John D.E. Gabrieli:** Conceptualization, Funding acquisition, Supervision, Writing - review & editing. **Joanna A. Christodoulou:** Conceptualization, Funding acquisition, Investigation, Supervision, Writing - review & editing.

## Acknowledgments

We thank all the participants and their families for volunteering their time to participate in the study. We also thank Atsushi Takahashi, Sheeba Arnold Anteraper, and Steven Shannon at the Athinoula A. Martinos Imaging Center at the McGovern Institute for Brain Research, Massachusetts Institute of Technology for technical assistance. We thank our colleagues and team members Rachel Romeo, Patricia Chang, Abigail Cyr, Pamela Hook, Jiayi Lin, Jack Murtagh, and Carly Schimmel.

## Supplementary Materials

**Figure S1:**
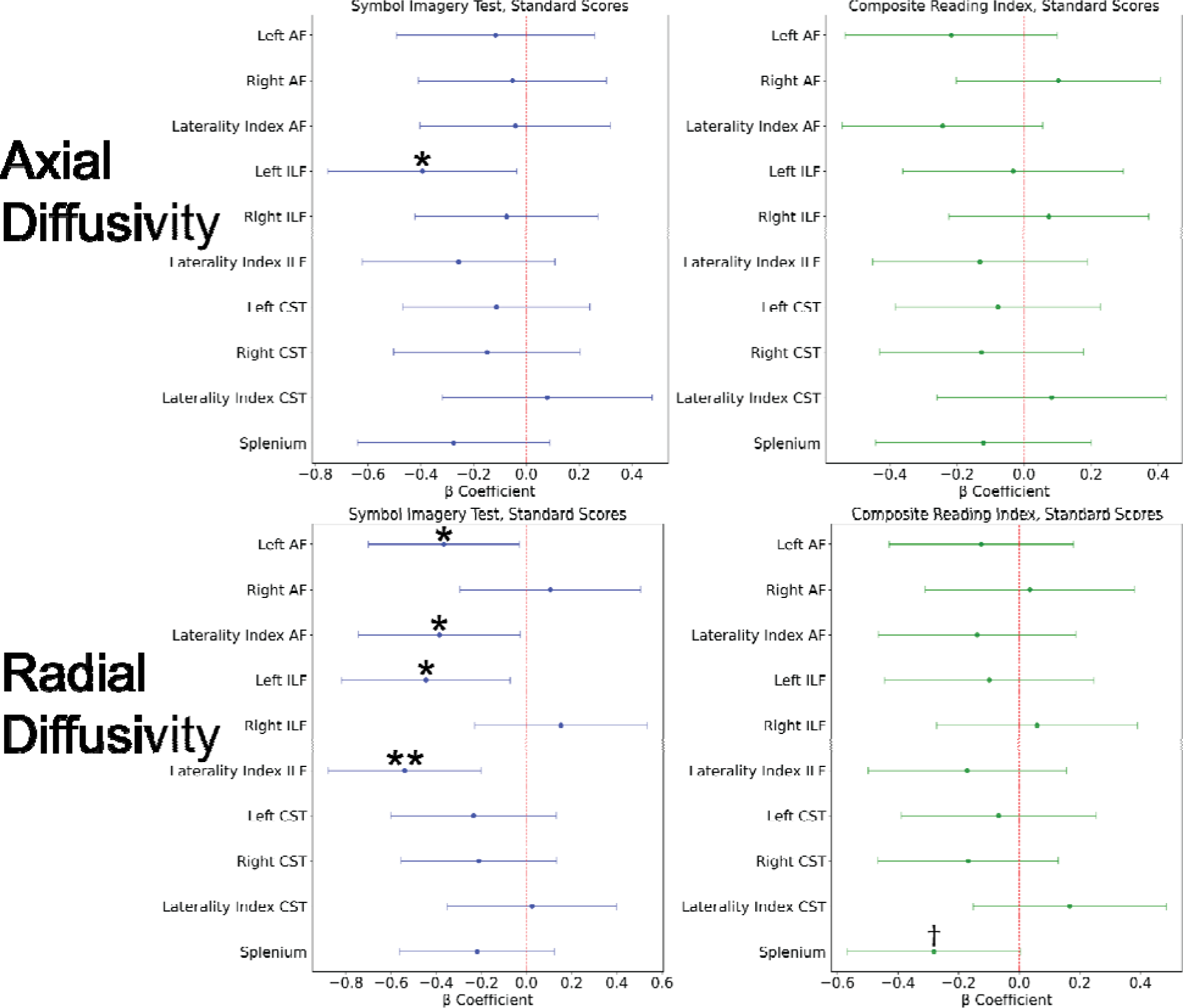
β coefficient plots for each model using axial diffusivity (top) and radial diffusivity (bottom). β coefficients quantify the relationship between the reading predictor of interest and the microstructural metric, after accounting for nuisance regressors in the models (age, sex, motion index). Error bars represent the 95% confidence interval surrounding the coefficient. †: p < 0.1, *: p < 0.05, **: p_FDR_ < 0.05. Abbreviations: AF - Arcuate Fasciculus; ILF - Inferior Longitudinal Fasciculus; CST - Corticospinal Tract.

**Figure S2:**
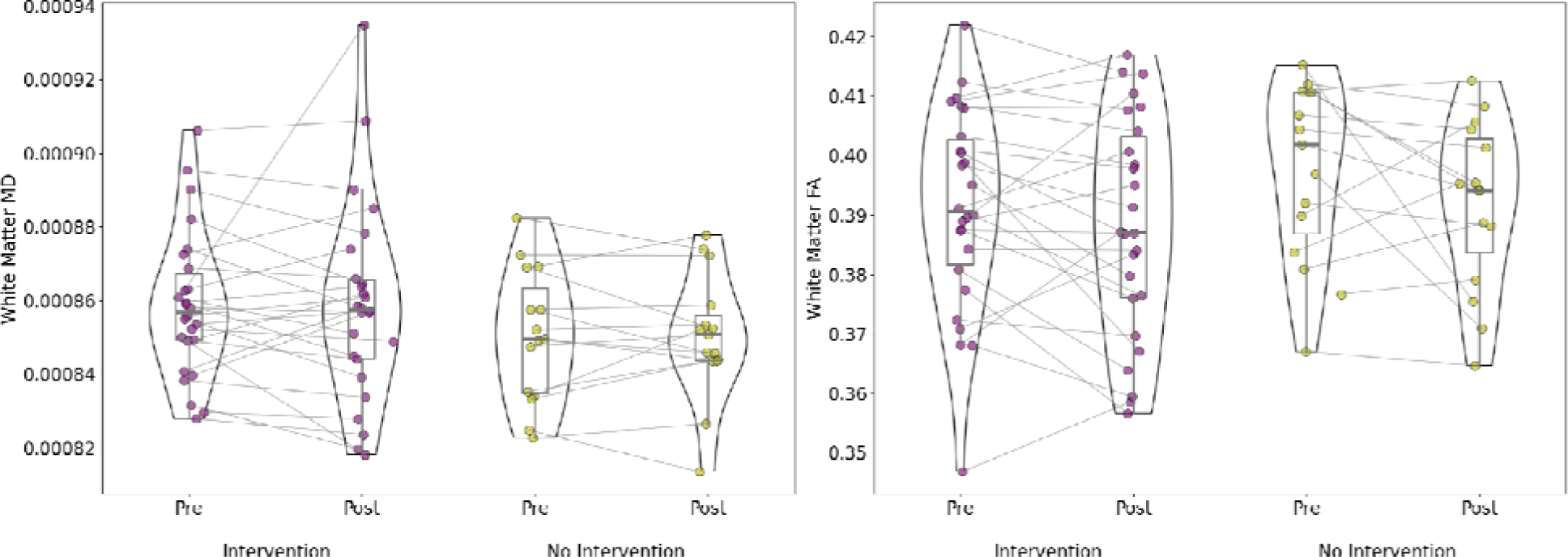
Changes in white matter average mean diffusivity (MD; left) and Fractional Anisotropy (FA; right) for intervention (purple) and non-intervention (yellow) participants. Paired t-tests were used to compare pre and post scores within groups, and two-sample t-tests were used to compare scores at a given time point across groups. No comparisons were statistically significant at a threshold of p < 0.05.

**Figure S3:**
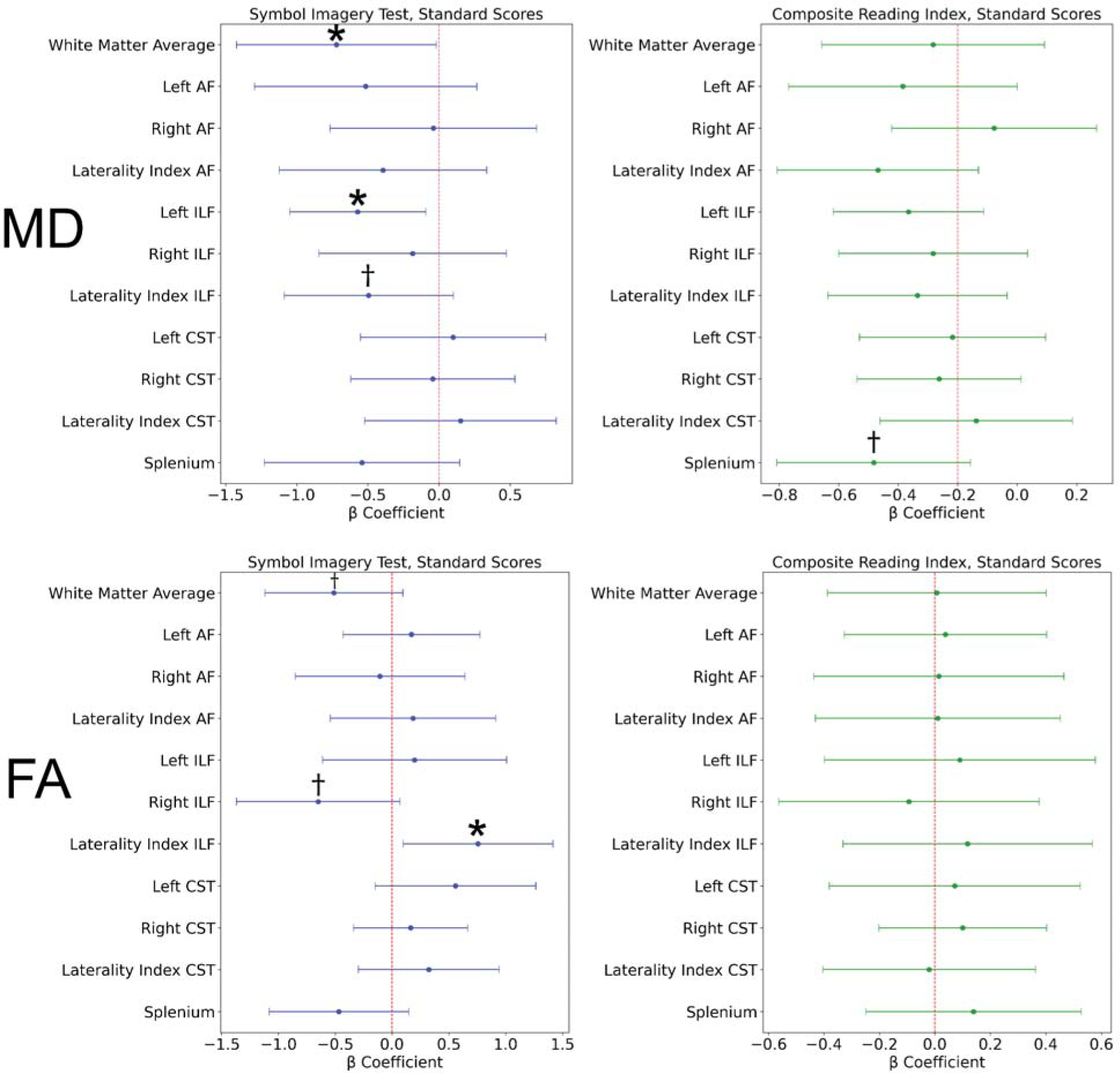
β coefficient plots for each model run only on the participants who completed the reading intervention (n = 26). β coefficients quantify the relationship between the reading predictor of interest and the microstructural metric, after accounting for nuisance regressors in the models (age, sex, motion index). Error bars represent the 95% confidence interval surrounding the coefficient. †: p < 0.1, *: p < 0.05, **: p_FDR_ < 0.05. Abbreviations: AF - Arcuate Fasciculus; ILF - Inferior Longitudinal Fasciculus; CST - Corticospinal Tract.

### Cross-Sectional Relationships Between White Matter Microstructure and Reading Scores

We ran models to look for relationships between white matter microstructure and reading performance cross-sectionally at each time point across all participants (*Figure S4*). Nuisance regressors included sex, age, and motion index specific to the time point. Across all participants at the beginning of the summer, decreased leftward ILF MD laterality (*p* < 0.05, ΔR^2^ = 0.085), and both lower MD (*p* < 0.1, ΔR^2^ = 0.058) and higher FA (*p* < 0.1, ΔR^2^ = 0.064) in the left ILF were associated with better SIT scores. Additionally, higher FA in the right ILF (*p* < 0.05, ΔR^2^_adj_ = 0.106) and right CST (*p* < 0.05, ΔR^2^ = 0.069), as well as lower MD in the right CST (*p* < 0.05, ΔR^2^ = 0.079), were related to better initial composite reading index scores. At the end of the summer, better composite reading index scores were associated with higher FA (*p* < 0.05, ΔR^2^ = 0.084) and lower MD (*p* < 0.05, ΔR^2^ = 0.086), in the right CST. No tract microstructure values were associated with SIT scores at the end of the summer. Decreasing rightward laterality in right CST MD was associated with better composite reading index scores at the end of the summer (*p* = 0.001, *p_FDR_*< 0.05; ΔR^2^ = 0.209). This was the only test that survived multiple comparison correction in the cross-sectional analyses.

**Figure S4:**
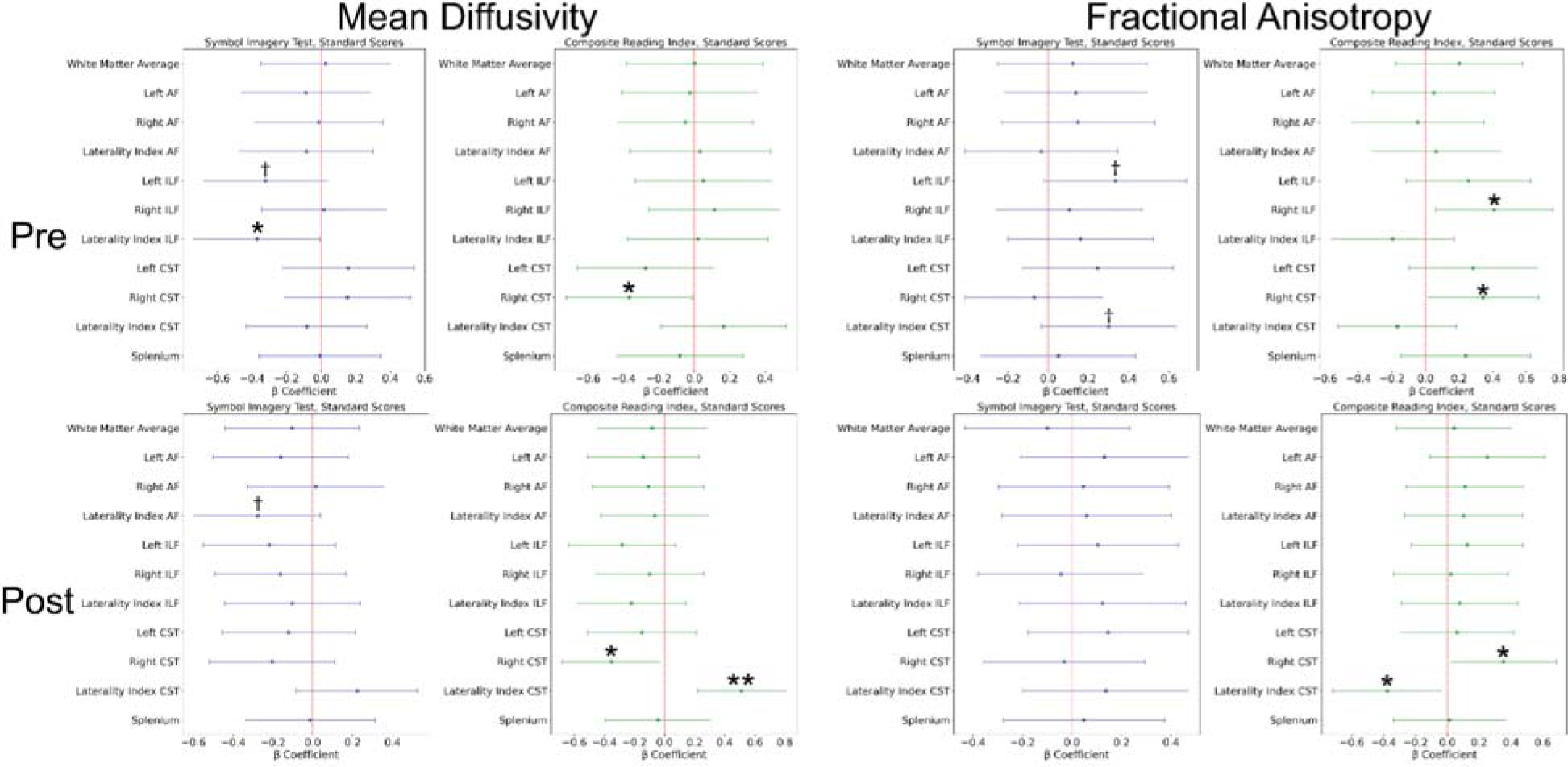
β coefficient plots for each model in the cross-sectional analyses. β coefficients quantify the relationship between the reading predictor of interest and the microstructural metric, after accounting for nuisance regressors in the models (age, sex, motion index). Error bars represent the 95% confidence interval surrounding the coefficient. †: p < 0.1, *: p < 0.05, **: p_FDR_ < 0.05. Abbreviations: AF - Arcuate Fasciculus; ILF - Inferior Longitudinal Fasciculus; CST - Corticospinal Tract.

## References

Andersson, J. L. R., & Sotiropoulos, S. N. (2016). An integrated approach to correction for off-resonance effects and subject movement in diffusion MR imaging. NeuroImage, 125, 1063–1078. 10.1016/j.neuroimage.2015.10.019

Badian, N. A. (2001). Phonological and orthographic processing: Their roles in reading prediction. Annals of Dyslexia, 51(1), 177–202. 10.1007/s11881-001-0010-5

Banfi, C., Koschutnig, K., Moll, K., Schulte-Körne, G., Fink, A., & Landerl, K. (2018). White matter alterations and tract lateralization in children with dyslexia and isolated spelling deficits. Human Brain Mapping, 40(3), 765–776. 10.1002/hbm.24410

Barquero, L. A., Davis, N., & Cutting, L. E. (2014). Neuroimaging of reading intervention: A systematic review and activation likelihood estimate meta-analysis. PloS One, 9(1), e83668.

Basser, P. J., Mattiello, J., & LeBihan, D. (1994). MR diffusion tensor spectroscopy and imaging. Biophysical Journal, 66(1), 259–267. 10.1016/S0006-3495(94)80775-1

Beaulieu, C. (2002). The basis of anisotropic water diffusion in the nervous system – a technical review. NMR in Biomedicine, 15(7–8), 435–455. 10.1002/nbm.782

Behler, A., Kassubek, J., & Müller, H.-P. (2021). Age-Related Alterations in DTI Metrics in the Human Brain—Consequences for Age Correction. Frontiers in Aging Neuroscience, 13. https://www.frontiersin.org/articles/10.3389/fnagi.2021.682109

Behrens, T. E. J., Berg, H. J., Jbabdi, S., Rushworth, M. F. S., & Woolrich, M. W. (2007). Probabilistic diffusion tractography with multiple fibre orientations: What can we gain? NeuroImage, 34(1), 144–155. 10.1016/j.neuroimage.2006.09.018

Bell, N. (1997). Seeing stars: Symbol imagery for phonemic awareness, sight words and spelling. Gander Publishing Avila Beach, CA.

Bell, N. (2010). Symbol imagery test. Gander Publishing.

Benjamini, Y., & Hochberg, Y. (1995). Controlling the False Discovery Rate: A Practical and Powerful Approach to Multiple Testing. Journal of the Royal Statistical Society: Series B (Methodological), 57(1), 289–300. 10.1111/j.2517-6161.1995.tb02031.x

Ben-Shachar, M., Dougherty, R. F., & Wandell, B. A. (2007). White matter pathways in reading. Current Opinion in Neurobiology, 17(2), 258–270. 10.1016/j.conb.2007.03.006

Blumenfeld-Katzir, T., Pasternak, O., Dagan, M., & Assaf, Y. (2011). Diffusion MRI of Structural Brain Plasticity Induced by a Learning and Memory Task. PLOS ONE, 6(6), e20678. 10.1371/journal.pone.0020678

Borchers, L. R., Bruckert, L., Dodson, C. K., Travis, K. E., Marchman, V. A., Ben-Shachar, M., & Feldman, H. M. (2019). Microstructural properties of white matter pathways in relation to subsequent reading abilities in children: A longitudinal analysis. Brain Structure & Function, 224(2), 891–905. 10.1007/s00429-018-1813-z

Bouhali, F., de Schotten, M. T., Pinel, P., Poupon, C., Mangin, J.-F., Dehaene, S., & Cohen, L. (2014). Anatomical connections of the visual word form area. Journal of Neuroscience, 34(46), 15402–15414.

Braid, J., & Richlan, F. (2022). The Functional Neuroanatomy of Reading Intervention. Frontiers in Neuroscience, 16, 921931. 10.3389/fnins.2022.921931

Catani, M., Jones, D. K., & Ffytche, D. H. (2005). Perisylvian language networks of the human brain. Annals of Neurology, 57(1), 8–16. 10.1002/ana.20319

Catani, M., & Mesulam, M. (2008). The arcuate fasciculus and the disconnection theme in language and aphasia: History and current state. Cortex, 44(8), 953–961. 10.1016/j.cortex.2008.04.002

Catts, H. W. (1993). The Relationship Between Speech-Language Impairments and Reading Disabilities. Journal of Speech, Language, and Hearing Research, 36(5), 948–958. 10.1044/jshr.3605.948

Christodoulou, J. A., Cyr, A., Murtagh, J., Chang, P., Lin, J., Guarino, A. J., Hook, P., & Gabrieli, J. D. E. (2017). Impact of Intensive Summer Reading Intervention for Children With Reading Disabilities and Difficulties in Early Elementary School. Journal of Learning Disabilities, 50(2), 115–127. 10.1177/0022219415617163

Connally, E. L., Ward, D., Howell, P., & Watkins, K. E. (2014). Disrupted white matter in language and motor tracts in developmental stuttering. Brain and Language, 131, 25–35. 10.1016/j.bandl.2013.05.013

Cooper, H., Nye, B., Charlton, K., Lindsay, J., & Greathouse, S. (1996). The Effects of Summer Vacation on Achievement Test Scores: A Narrative and Meta-Analytic Review. Review of Educational Research, 66(3), 227–268. 10.3102/00346543066003227

Cui, Z., Xia, Z., Su, M., Shu, H., & Gong, G. (2016). Disrupted white matter connectivity underlying developmental dyslexia: A machine learning approach. Human Brain Mapping, 37(4), 1443–1458. 10.1002/hbm.23112

Davis, B. R., Garza, A., & Church, J. A. (2022). Key considerations for child and adolescent MRI data collection. Frontiers in Neuroimaging, 1, 981947. 10.3389/fnimg.2022.981947

Davison, K. E., Zuk, J., Mullin, L. J., Ozernov-Palchik, O., Norton, E., Gabrieli, J. D. E., Yu, X., & Gaab, N. (2022). Examining Shared Reading and White Matter Organization in Kindergarten in Relation to Subsequent Language and Reading Abilities: A Longitudinal Investigation. Journal of Cognitive Neuroscience, 1–17. 10.1162/jocn_a_01944

de Bie, H. M. A., Boersma, M., Wattjes, M. P., Adriaanse, S., Vermeulen, R. J., Oostrom, K. J., Huisman, J., Veltman, D. J., & Delemarre-Van de Waal, H. A. (2010). Preparing children with a mock scanner training protocol results in high quality structural and functional MRI scans. European Journal of Pediatrics, 169(9), 1079–1085. 10.1007/s00431-010-1181-z

De Santis, S., Drakesmith, M., Bells, S., Assaf, Y., & Jones, D. K. (2014). Why diffusion tensor MRI does well only some of the time: Variance and covariance of white matter tissue microstructure attributes in the living human brain. NeuroImage, 89, 35–44. 10.1016/j.neuroimage.2013.12.003

Dehaene, S., & Cohen, L. (2011). The unique role of the visual word form area in reading. Trends in Cognitive Sciences, 15(6), 254–262. 10.1016/j.tics.2011.04.003

Donnelly, P. M., Huber, E., & Yeatman, J. D. (2019). Intensive Summer Intervention Drives Linear Growth of Reading Skill in Struggling Readers. Frontiers in Psychology, 10, 1900. 10.3389/fpsyg.2019.01900

Duffau, H., Herbet, G., & Moritz-Gasser, S. (2013). Toward a pluri-component, multimodal, and dynamic organization of the ventral semantic stream in humans: Lessons from stimulation mapping in awake patients. Frontiers in Systems Neuroscience, 7. https://www.frontiersin.org/articles/10.3389/fnsys.2013.00044

Economou, M., Bempt, F. V., Van Herck, S., Wouters, J., Ghesquière, P., Vanderauwera, J., & Vandermosten, M. (2023). Myelin plasticity during early literacy training in at-risk pre-readers. Cortex, 167, 86–100. 10.1016/j.cortex.2023.05.023

Economou, M., Van Herck, S., Vanden Bempt, F., Glatz, T., Wouters, J., Ghesquière, P., Vanderauwera, J., & Vandermosten, M. (2022). Investigating the impact of early literacy training on white matter structure in prereaders at risk for dyslexia. Cerebral Cortex, 32(21), 4684–4697. 10.1093/cercor/bhab510

Eden, G. F., Jones, K. M., Cappell, K., Gareau, L., Wood, F. B., Zeffiro, T. A., Dietz, N. A. E., Agnew, J. A., & Flowers, D. L. (2004). Neural Changes following Remediation in Adult Developmental Dyslexia. Neuron, 44(3), 411–422. 10.1016/j.neuron.2004.10.019

Enge, A., Friederici, A. D., & Skeide, M. A. (2020). A meta-analysis of fMRI studies of language comprehension in children. NeuroImage, 215, 116858. 10.1016/j.neuroimage.2020.116858

Entwisle, D. R., Alexander, K. L., & Olson, L. S. (1997). Children, schools, and inequality. Routledge.

Epelbaum, S., Pinel, P., Gaillard, R., Delmaire, C., Perrin, M., Dupont, S., Dehaene, S., & Cohen, L. (2008). Pure alexia as a disconnection syndrome: New diffusion imaging evidence for an old concept. Cortex; a Journal Devoted to the Study of the Nervous System and Behavior, 44(8), 962–974. 10.1016/j.cortex.2008.05.003

Faul, F., Erdfelder, E., Buchner, A., & Lang, A.-G. (2009). Statistical power analyses using G* Power 3.1: Tests for correlation and regression analyses. Behavior Research Methods, 41(4), 1149–1160.

Fedorenko, E., Behr, M. K., & Kanwisher, N. (2011). Functional specificity for high-level linguistic processing in the human brain. Proceedings of the National Academy of Sciences, 108(39), 16428–16433. 10.1073/pnas.1112937108

Fields, R. D. (2015). A new mechanism of nervous system plasticity: Activity-dependent myelination. Nature Reviews Neuroscience, 16(12), Article 12. 10.1038/nrn4023

Fields, R. D., Araque, A., Johansen-Berg, H., Lim, S.-S., Lynch, G., Nave, K.-A., Nedergaard, M., Perez, R., Sejnowski, T., & Wake, H. (2014). Glial Biology in Learning and Cognition. The Neuroscientist, 20(5), 426–431. 10.1177/1073858413504465

Fischl, B. (2012). FreeSurfer. NeuroImage, 62(2), 774–781. 10.1016/j.neuroimage.2012.01.021

Fischl, B., Salat, D. H., Busa, E., Albert, M., Dieterich, M., Haselgrove, C., van der Kouwe, A., Killiany, R., Kennedy, D., Klaveness, S., Montillo, A., Makris, N., Rosen, B., & Dale, A. M. (2002). Whole Brain Segmentation: Automated Labeling of Neuroanatomical Structures in the Human Brain. Neuron, 33(3), 341–355. 10.1016/S0896-6273(02)00569-X

Friedrich, P., Fraenz, C., Schlüter, C., Ocklenburg, S., Mädler, B., Güntürkün, O., & Genç, E. (2020). The Relationship Between Axon Density, Myelination, and Fractional Anisotropy in the Human Corpus Callosum. Cerebral Cortex (New York, N.Y.: 1991), 30(4), 2042–2056. 10.1093/cercor/bhz221

Gao, P., Wang, Y.-S., Lu, Q.-Y., Rong, M.-J., Fan, X.-R., Holmes, A. J., Dong, H.-M., Li, H.-F., & Zuo, X.-N. (2023). Brief mock-scan training reduces head motion during real scanning for children: A growth curve study. Developmental Cognitive Neuroscience, 61, 101244. 10.1016/j.dcn.2023.101244

Genc, S., Malpas, C. B., Holland, S. K., Beare, R., & Silk, T. J. (2017). Neurite density index is sensitive to age related differences in the developing brain. NeuroImage, 148, 373–380. 10.1016/j.neuroimage.2017.01.023

Genc, S., Tax, C. M. W., Raven, E. P., Chamberland, M., Parker, G. D., & Jones, D. K. (2020). Impact of b-value on estimates of apparent fibre density. Human Brain Mapping, 41(10), 2583–2595. 10.1002/hbm.24964

Good, R. H., Gruba, J., & Kaminski, R. A. (2002). Best Practices in Using Dynamic Indicators of Basic Early Literacy Skills (DIBELS) in an Outcomes-Driven Model. In Best practices in school psychology IV, Vols. 1-2 (pp. 699–720). National Association of School Psychologists.

Greene, D. J., Koller, J. M., Hampton, J. M., Wesevich, V., Van, A. N., Nguyen, A. L., Hoyt, C. R., McIntyre, L., Earl, E. A., & Klein, R. L. (2018). Behavioral interventions for reducing head motion during MRI scans in children. Neuroimage, 171, 234–245.

Greve, D. N., & Fischl, B. (2009). Accurate and robust brain image alignment using boundary-based registration. Neuroimage, 48(1), 63–72.

Gullick, M. M., & Booth, J. R. (2015). The direct segment of the arcuate fasciculus is predictive of longitudinal reading change. Developmental Cognitive Neuroscience, 13, 68–74. 10.1016/j.dcn.2015.05.002

Han, Z., Ma, Y., Gong, G., Huang, R., Song, L., & Bi, Y. (2016). White matter pathway supporting phonological encoding in speech production: A multi-modal imaging study of brain damage patients. Brain Structure and Function, 221(1), 577–589. 10.1007/s00429-014-0926-2

Hayiou-Thomas Marianna E., Harlaar, N., Dale, P. S., & Plomin, R. (2010). Preschool Speech, Language Skills, and Reading at 7, 9, and 10 Years: Etiology of the Relationship. Journal of Speech, Language, and Hearing Research, 53(2), 311–332. 10.1044/1092-4388(2009/07-0145)

Herbet, G., Zemmoura, I., & Duffau, H. (2018). Functional Anatomy of the Inferior Longitudinal Fasciculus: From Historical Reports to Current Hypotheses. Frontiers in Neuroanatomy, 12. https://www.frontiersin.org/articles/10.3389/fnana.2018.00077

Hernández, M., Guerrero, G. D., Cecilia, J. M., García, J. M., Inuggi, A., Jbabdi, S., Behrens, T. E. J., & Sotiropoulos, S. N. (2013). Accelerating Fibre Orientation Estimation from Diffusion Weighted Magnetic Resonance Imaging Using GPUs. PLOS ONE, 8(4), e61892. 10.1371/journal.pone.0061892

Hoeft, F., McCandliss, B. D., Black, J. M., Gantman, A., Zakerani, N., Hulme, C., Lyytinen, H., Whitfield-Gabrieli, S., Glover, G. H., & Reiss, A. L. (2011). Neural systems predicting long-term outcome in dyslexia. Proceedings of the National Academy of Sciences, 108(1), 361–366.

Huber, E., Donnelly, P. M., Rokem, A., & Yeatman, J. D. (2018). Rapid and widespread white matter plasticity during an intensive reading intervention. Nature Communications, 9(1), 1–13.

Huber, E., Mezer, A., & Yeatman, J. D. (2021). Neurobiological underpinnings of rapid white matter plasticity during intensive reading instruction. NeuroImage, 243, 118453. 10.1016/j.neuroimage.2021.118453

Jang, H., Lee, J. Y., Lee, K. I., & Park, K. M. (2017). Are there differences in brain morphology according to handedness? Brain and Behavior, 7(7), e00730. 10.1002/brb3.730

Jbabdi, S., Sotiropoulos, S. N., Savio, A. M., Graña, M., & Behrens, T. E. J. (2012). Model-based analysis of multishell diffusion MR data for tractography: How to get over fitting problems. Magnetic Resonance in Medicine, 68(6), 1846–1855. 10.1002/mrm.24204

Jelescu, I. O., Palombo, M., Bagnato, F., & Schilling, K. G. (2020). Challenges for biophysical modeling of microstructure. Journal of Neuroscience Methods, 344, 108861. 10.1016/j.jneumeth.2020.108861

Jeurissen, B., Leemans, A., Tournier, J.-D., Jones, D. K., & Sijbers, J. (2013). Investigating the prevalence of complex fiber configurations in white matter tissue with diffusion magnetic resonance imaging. Human Brain Mapping, 34(11), 2747–2766.

Jones, D. K., Knösche, T. R., & Turner, R. (2013). White matter integrity, fiber count, and other fallacies: The do’s and don’ts of diffusion MRI. NeuroImage, 73, 239–254. 10.1016/j.neuroimage.2012.06.081

Kaufman, A. S., Kaufman, Nadeen L. (2004). Kaufman brief intelligence test KBIT 2LJ; manual. /z-wcorg/.

Keller, T. A., & Just, M. A. (2009). Altering Cortical Connectivity: Remediation-Induced Changes in the White Matter of Poor Readers. Neuron, 64(5), 624–631. 10.1016/j.neuron.2009.10.018

King, K. M., Littlefield, A. K., McCabe, C. J., Mills, K. L., Flournoy, J., & Chassin, L. (2018). Longitudinal modeling in developmental neuroimaging research: Common challenges, and solutions from developmental psychology. Developmental Cognitive Neuroscience, 33, 54–72. 10.1016/j.dcn.2017.11.009

Koirala, N., Perdue, M. V., Su, X., Grigorenko, E. L., & Landi, N. (2021). Neurite density and arborization is associated with reading skill and phonological processing in children. NeuroImage, 241, 118426. 10.1016/j.neuroimage.2021.118426

Kronfeld-Duenias, V., Amir, O., Ezrati-Vinacour, R., Civier, O., & Ben-Shachar, M. (2016). The frontal aslant tract underlies speech fluency in persistent developmental stuttering. Brain Structure and Function, 221(1), 365–381. 10.1007/s00429-014-0912-8

Kuhfeld, M., Lewis, K., & Peltier, T. (2023). Reading achievement declines during the COVID-19 pandemic: Evidence from 5 million U.S. students in grades 3–8. Reading and Writing, 36(2), 245–261. 10.1007/s11145-022-10345-8

Kurtzer, G. M., Sochat, V., & Bauer, M. W. (2017). Singularity: Scientific containers for mobility of compute. PLOS ONE, 12(5), e0177459. 10.1371/journal.pone.0177459

Langer, N., Peysakhovich, B., Zuk, J., Drottar, M., Sliva, D. D., Smith, S., Becker, B. L. C., Grant, P. E., & Gaab, N. (2017). White Matter Alterations in Infants at Risk for Developmental Dyslexia. Cerebral Cortex, 27(2), 1027–1036. 10.1093/cercor/bhv281

Leemans, A., & Jones, D. K. (2009). The B-matrix must be rotated when correcting for subject motion in DTI data. Magnetic Resonance in Medicine, 61(6), 1336–1349. 10.1002/mrm.21890

Lerma-Usabiaga, G., Carreiras, M., & Paz-Alonso, P. M. (2018). Converging evidence for functional and structural segregation within the left ventral occipitotemporal cortex in reading. Proceedings of the National Academy of Sciences, 115(42), E9981–E9990. 10.1073/pnas.1803003115

López-Vicente, M., Lamballais, S., Louwen, S., Hillegers, M., Tiemeier, H., Muetzel, R. L., & White, T. (2021). White matter microstructure correlates of age, sex, handedness and motor ability in a population-based sample of 3031 school-age children. NeuroImage, 227, 117643. 10.1016/j.neuroimage.2020.117643

Maffei, C., Lee, C., Planich, M., Ramprasad, M., Ravi, N., Trainor, D., Urban, Z., Kim, M., Jones, R. J., Henin, A., Hofmann, S. G., Pizzagalli, D. A., Auerbach, R. P., Gabrieli, J. D. E., Whitfield-Gabrieli, S., Greve, D. N., Haber, S. N., & Yendiki, A. (2021). Using diffusion MRI data acquired with ultra-high gradient strength to improve tractography in routine-quality data. NeuroImage, 245, 118706. 10.1016/j.neuroimage.2021.118706

McKinney, W. (2011). pandas: A foundational Python library for data analysis and statistics. Python for High Performance and Scientific Computing, 14(9), 1–9.

Meisler, S. L., & Gabrieli, J. D. (2022a). A Large-Scale Investigation of White Matter Microstructural Associations with Reading Ability. NeuroImage, 118909.

Meisler, S. L., & Gabrieli, J. D. (2022b). Fiber-specific structural properties relate to reading skills in children and adolescents. eLife, 11, e82088. 10.7554/eLife.82088

Metzler-Baddeley, C., Foley, S., de Santis, S., Charron, C., Hampshire, A., Caeyenberghs, K., & Jones, D. K. (2017). Dynamics of White Matter Plasticity Underlying Working Memory Training: Multimodal Evidence from Diffusion MRI and Relaxometry. Journal of Cognitive Neuroscience, 29(9), 1509–1520. 10.1162/jocn_a_01127

Moreau, D., Stonyer, J. E., McKay, N. S., & Waldie, K. E. (2018). No evidence for systematic white matter correlates of dyslexia: An Activation Likelihood Estimation meta-analysis. Brain Research, 1683, 36–47. 10.1016/j.brainres.2018.01.014

Murphy, K. A., Jogia, J., & Talcott, J. B. (2019). On the neural basis of word reading: A meta-analysis of fMRI evidence using activation likelihood estimation. Journal of Neurolinguistics, 49, 71–83.

Myers, C. A., Vandermosten, M., Farris, E. A., Hancock, R., Gimenez, P., Black, J. M., Casto, B., Drahos, M., Tumber, M., Hendren, R. L., Hulme, C., & Hoeft, F. (2014). White Matter Morphometric Changes Uniquely Predict Children’s Reading Acquisition. Psychological Science, 25(10), 1870–1883. 10.1177/0956797614544511

Ng, S., Moritz-Gasser, S., Lemaitre, A.-L., Duffau, H., & Herbet, G. (2021). White matter disconnectivity fingerprints causally linked to dissociated forms of alexia. Communications Biology, 4(1), Article 1. 10.1038/s42003-021-02943-z

Northam, G. B., Liégeois, F., Chong, W. K., Baker, K., Tournier, J.-D., Wyatt, J. S., Baldeweg, T., & Morgan, A. (2012). Speech and Oromotor Outcome in Adolescents Born Preterm: Relationship to Motor Tract Integrity. The Journal of Pediatrics, 160(3), 402–408.e1. 10.1016/j.jpeds.2011.08.055

Northam, G. B., Morgan, A. T., Fitzsimmons, S., Baldeweg, T., & Liégeois, F. J. (2019). Corticobulbar Tract Injury, Oromotor Impairment and Language Plasticity in Adolescents Born Preterm. Frontiers in Human Neuroscience, 13. https://www.frontiersin.org/articles/10.3389/fnhum.2019.00045

Partanen, M., Kim, D. H., Rauscher, A., Siegel, L. S., & Giaschi, D. E. (2021). White matter but not grey matter predicts change in reading skills after intervention. Dyslexia, 27(2), 224–244.

Perdue, M. V., Mahaffy, K., Vlahcevic, K., Wolfman, E., Erbeli, F., Richlan, F., & Landi, N. (2022). Reading intervention and neuroplasticity: A systematic review and meta-analysis of brain changes associated with reading intervention. Neuroscience & Biobehavioral Reviews, 132, 465–494.

Raffelt, D. A., Tournier, J.-D., Smith, R. E., Vaughan, D. N., Jackson, G., Ridgway, G. R., & Connelly, A. (2017). Investigating white matter fibre density and morphology using fixel-based analysis. NeuroImage, 144, 58–73. 10.1016/j.neuroimage.2016.09.029

Reuter, M., & Fischl, B. (2011). Avoiding asymmetry-induced bias in longitudinal image processing. NeuroImage, 57(1), 19–21. 10.1016/j.neuroimage.2011.02.076

Reuter, M., Rosas, H. D., & Fischl, B. (2010). Highly accurate inverse consistent registration: A robust approach. NeuroImage, 53(4), 1181–1196. 10.1016/j.neuroimage.2010.07.020

Reuter, M., Schmansky, N. J., Rosas, H. D., & Fischl, B. (2012). Within-subject template estimation for unbiased longitudinal image analysis. NeuroImage, 61(4), 1402–1418. 10.1016/j.neuroimage.2012.02.084

Richards, T. L., Berninger, V. W., Yagle, K. J., Abbott, R. D., & Peterson, D. J. (2017). Changes in DTI Diffusivity and fMRI Connectivity Cluster Coefficients for Students with and without Specific Learning Disabilities In Written Language: Brain’s Response to Writing Instruction. Journal of Nature and Science, 3(4), e350.

Romeo, R. R., Christodoulou, J. A., Halverson, K. K., Murtagh, J., Cyr, A. B., Schimmel, C., Chang, P., Hook, P. E., & Gabrieli, J. D. E. (2018). Socioeconomic Status and Reading Disability: Neuroanatomy and Plasticity in Response to Intervention. Cerebral Cortex, 28(7), 2297–2312. 10.1093/cercor/bhx131

Roy, E., Richie-Halford, A., Kruper, J., Narayan, M., Bloom, D., Nedelec, P., Rauschecker, A. M., Sugrue, L. P., Brown, T. T., Jernigan, T. L., McCandliss, B. D., Rokem, A., & Yeatman, J. D. (2024). White matter and literacy: A dynamic system in flux. Developmental Cognitive Neuroscience, 65, 101341. 10.1016/j.dcn.2024.101341

Sampaio-Baptista, C., & Johansen-Berg, H. (2017). White Matter Plasticity in the Adult Brain. Neuron, 96(6), 1239–1251. 10.1016/j.neuron.2017.11.026

Sampaio-Baptista, C., Khrapitchev, A. A., Foxley, S., Schlagheck, T., Scholz, J., Jbabdi, S., DeLuca, G. C., Miller, K. L., Taylor, A., Thomas, N., Kleim, J., Sibson, N. R., Bannerman, D., & Johansen-Berg, H. (2013). Motor Skill Learning Induces Changes in White Matter Microstructure and Myelination. Journal of Neuroscience, 33(50), 19499–19503. 10.1523/JNEUROSCI.3048-13.2013

Saur, D., Kreher, B. W., Schnell, S., Kümmerer, D., Kellmeyer, P., Vry, M.-S., Umarova, R., Musso, M., Glauche, V., Abel, S., Huber, W., Rijntjes, M., Hennig, J., & Weiller, C. (2008). Ventral and dorsal pathways for language. Proceedings of the National Academy of Sciences, 105(46), 18035–18040. 10.1073/pnas.0805234105

Saygin, Z. M., Norton, E. S., Osher, D. E., Beach, S. D., Cyr, A. B., Ozernov-Palchik, O., Yendiki, A., Fischl, B., Gaab, N., & Gabrieli, J. D. E. (2013). Tracking the Roots of Reading Ability: White Matter Volume and Integrity Correlate with Phonological Awareness in Prereading and Early-Reading Kindergarten Children. Journal of Neuroscience, 33(33), 13251–13258. 10.1523/JNEUROSCI.4383-12.2013

Schilling, K. G., Rheault, F., Petit, L., Hansen, C. B., Nath, V., Yeh, F.-C., Girard, G., Barakovic, M., Rafael-Patino, J., Yu, T., Fischi-Gomez, E., Pizzolato, M., Ocampo-Pineda, M., Schiavi, S., Canales-Rodríguez, E. J., Daducci, A., Granziera, C., Innocenti, G., Thiran, J.-P., … Descoteaux, M. (2021). Tractography dissection variability: What happens when 42 groups dissect 14 white matter bundles on the same dataset? NeuroImage, 243, 118502. 10.1016/j.neuroimage.2021.118502

Schilling, K. G., Tax, C. M. W., Rheault, F., Hansen, C., Yang, Q., Yeh, F.-C., Cai, L., Anderson, A. W., & Landman, B. A. (2021). Fiber tractography bundle segmentation depends on scanner effects, vendor effects, acquisition resolution, diffusion sampling scheme, diffusion sensitization, and bundle segmentation workflow. NeuroImage, 242, 118451. 10.1016/j.neuroimage.2021.118451

Schilling, K. G., Tax, C. M. W., Rheault, F., Landman, B. A., Anderson, A. W., Descoteaux, M., & Petit, L. (2022). Prevalence of white matter pathways coming into a single white matter voxel orientation: The bottleneck issue in tractography. Human Brain Mapping, 43(4), 1196–1213. 10.1002/hbm.25697

Scholz, J., Klein, M. C., Behrens, T. E. J., & Johansen-Berg, H. (2009). Training induces changes in white matter architecture. Nature Neuroscience, 12(11), 1370–1371. 10.1038/nn.2412

Seabold, S., & Perktold, J. (2010). Statsmodels: Econometric and statistical modeling with python. Proceedings of the 9th Python in Science Conference, 57(61), 10.25080.

Shaywitz, S. E. (1998). Dyslexia. New England Journal of Medicine, 338(5), 307–312. 10.1056/NEJM199801293380507

Shin, J., Rowley, J., Chowdhury, R., Jolicoeur, P., Klein, D., Grova, C., Rosa-Neto, P., & Kobayashi, E. (2019). Inferior Longitudinal Fasciculus’ Role in Visual Processing and Language Comprehension: A Combined MEG-DTI Study. Frontiers in Neuroscience, 13, 875. 10.3389/fnins.2019.00875

Sihvonen, A. J., Virtala, P., Thiede, A., Laasonen, M., & Kujala, T. (2021). Structural white matter connectometry of reading and dyslexia. NeuroImage, 241, 118411.

Thiebaut de Schotten, M., Cohen, L., Amemiya, E., Braga, L. W., & Dehaene, S. (2014). Learning to Read Improves the Structure of the Arcuate Fasciculus. Cerebral Cortex, 24(4), 989–995. 10.1093/cercor/bhs383

Thornton, A., & Lee, P. (2000). Publication bias in meta-analysis: Its causes and consequences. Journal of Clinical Epidemiology, 53(2), 207–216. 10.1016/S0895-4356(99)00161-4

Torgesen, J. K., Wagner, R. K., & Rashotte, C. A. (2012). Test of word reading efficiency–second edition (TOWRE-2). Austin, TX: Pro-Ed.

Torgeson, J. K., Wagner, R. K., & Rashotte, C. A. (1999). Test Review: Test of Word Reading Efficiency (TOWRE). Pro-ed.

Van Der Auwera, S., Vandermosten, M., Wouters, J., Ghesquière, P., & Vanderauwera, J. (2021). A three-time point longitudinal investigation of the arcuate fasciculus throughout reading acquisition in children developing dyslexia. NeuroImage, 237, 118087. 10.1016/j.neuroimage.2021.118087

Vanderauwera, J., Wouters, J., Vandermosten, M., & Ghesquière, P. (2017). Early dynamics of white matter deficits in children developing dyslexia. Developmental Cognitive Neuroscience, 27, 69–77. 10.1016/j.dcn.2017.08.003

Vandermosten, M., Boets, B., Wouters, J., & Ghesquière, P. (2012). A qualitative and quantitative review of diffusion tensor imaging studies in reading and dyslexia. Neuroscience & Biobehavioral Reviews, 36(6), 1532–1552. 10.1016/j.neubiorev.2012.04.002

Vandermosten, M., Vanderauwera, J., Theys, C., De Vos, A., Vanvooren, S., Sunaert, S., Wouters, J., & Ghesquière, P. (2015). A DTI tractography study in pre-readers at risk for dyslexia. Developmental Cognitive Neuroscience, 14, 8–15. 10.1016/j.dcn.2015.05.006

Vigneau, M., Beaucousin, V., Hervé, P. Y., Duffau, H., Crivello, F., Houdé, O., Mazoyer, B., & Tzourio-Mazoyer, N. (2006). Meta-analyzing left hemisphere language areas: Phonology, semantics, and sentence processing. NeuroImage, 30(4), 1414–1432. 10.1016/j.neuroimage.2005.11.002

von Hippel, P. T., Workman, J., & Downey, D. B. (2018). Inequality in Reading and Math Skills Forms Mainly before Kindergarten: A Replication, and Partial Correction, of “Are Schools the Great Equalizer?” Sociology of Education, 91(4), 323–357. 10.1177/0038040718801760

Wagner, R. K., Torgesen, J. K., Rashotte, C. A., & Pearson, N. A. (1999). Comprehensive test of phonological processing: CTOPP. Pro-ed Austin, TX.

Wagner, R. K., Zirps, F. A., Edwards, A. A., Wood, S. G., Joyner, R. E., Becker, B. J., Liu, G., & Beal, B. (2020). The prevalence of Dyslexia: A new approach to its estimation. Journal of Learning Disabilities, 53(5), 354–365.

Walton, M., Dewey, D., & Lebel, C. (2018). Brain white matter structure and language ability in preschool-aged children. Brain and Language, 176, 19–25. 10.1016/j.bandl.2017.10.008

Wang, Y., Mauer, M. V., Raney, T., Peysakhovich, B., Becker, B. L., Sliva, D. D., & Gaab, N. (2017). Development of tract-specific white matter pathways during early reading development in at-risk children and typical controls. Cerebral Cortex, 27(4), 2469–2485.

Waskom, M. L. (2021). Seaborn: Statistical data visualization. Journal of Open Source Software, 6(60), 3021.

Weiner, K. S., Yeatman, J. D., & Wandell, B. A. (2017). The posterior arcuate fasciculus and the vertical occipital fasciculus. Cortex; a Journal Devoted to the Study of the Nervous System and Behavior, 97, 274–276. 10.1016/j.cortex.2016.03.012

Winklewski, P. J., Sabisz, A., Naumczyk, P., Jodzio, K., Szurowska, E., & Szarmach, A. (2018). Understanding the Physiopathology Behind Axial and Radial Diffusivity Changes—What Do We Know? Frontiers in Neurology, 9. 10.3389/fneur.2018.00092

Wolf, M., & Denckla, M. B. (2005). RAN/RAS: Rapid automatized naming and rapid alternating stimulus tests. Pro-ed Austin, TX.

Woodcock, R. W. (2011). Woodcock reading mastery tests: WRMT-III. Pearson.

Wu, L.-Y., Xu, Y., Chen, L.-L., Yang, W.-R., Li, Y., Shang, S.-A., Luo, X.-F., Xia, W., Xia, J., & Zhang, H.-Y. (2022). Test-retest reliability of diffusion kurtosis imaging metrics in the healthy adult brain. Neuroimage: Reports, 2(3), 100098. 10.1016/j.ynirp.2022.100098

Xin, W., & Chan, J. R. (2020). Myelin plasticity: Sculpting circuits in learning and memory. Nature Reviews Neuroscience, 21(12), Article 12. 10.1038/s41583-020-00379-8

Yeatman, J. D., Dougherty, R. F., Ben-Shachar, M., & Wandell, B. A. (2012). Development of white matter and reading skills. Proceedings of the National Academy of Sciences, 109(44), E3045–E3053.

Yeatman, J. D., Rauschecker, A. M., & Wandell, B. A. (2013). Anatomy of the visual word form area: Adjacent cortical circuits and long-range white matter connections. Brain and Language, 125(2), 146–155. 10.1016/j.bandl.2012.04.010

Yeatman, J. D., & White, A. L. (2021). Reading: The Confluence of Vision and Language. Annual Review of Vision Science, 7(1), 487–517. 10.1146/annurev-vision-093019-113509

Yeh, F.-C., Verstynen, T. D., Wang, Y., Fernández-Miranda, J. C., & Tseng, W.-Y. I. (2013). Deterministic Diffusion Fiber Tracking Improved by Quantitative Anisotropy. PLOS ONE, 8(11), e80713. 10.1371/journal.pone.0080713

Yendiki, A., Koldewyn, K., Kakunoori, S., Kanwisher, N., & Fischl, B. (2014). Spurious group differences due to head motion in a diffusion MRI study. NeuroImage, 88, 79–90. 10.1016/j.neuroimage.2013.11.027

Yendiki, A., Panneck, P., Srinivasan, P., Stevens, A., Zöllei, L., Augustinack, J., Wang, R., Salat, D., Ehrlich, S., Behrens, T., Jbabdi, S., Gollub, R., & Fischl, B. (2011). Automated Probabilistic Reconstruction of White-Matter Pathways in Health and Disease Using an Atlas of the Underlying Anatomy. Frontiers in Neuroinformatics, 5. https://www.frontiersin.org/articles/10.3389/fninf.2011.00023

Yendiki, A., Reuter, M., Wilkens, P., Rosas, H. D., & Fischl, B. (2016). Joint reconstruction of white-matter pathways from longitudinal diffusion MRI data with anatomical priors. NeuroImage, 127, 277–286. 10.1016/j.neuroimage.2015.12.003

Yu, Q., Peng, Y., Kang, H., Peng, Q., Ouyang, M., Slinger, M., Hu, D., Shou, H., Fang, F., & Huang, H. (2020). Differential White Matter Maturation from Birth to 8 Years of Age. Cerebral Cortex, 30(4), 2674–2690. 10.1093/cercor/bhz268

Zemmoura, I., Herbet, G., Moritz-Gasser, S., & Duffau, H. (2015). New insights into the neural network mediating reading processes provided by cortico-subcortical electrical mapping. Human Brain Mapping, 36(6), 2215–2230. 10.1002/hbm.22766

Zhang, H., Schneider, T., Wheeler-Kingshott, C. A., & Alexander, D. C. (2012). NODDI: Practical in vivo neurite orientation dispersion and density imaging of the human brain. NeuroImage, 61(4), 1000–1016. 10.1016/j.neuroimage.2012.03.072

Zuk, J., Dunstan, J., Norton, E., Yu, X., Ozernov-Palchik, O., Wang, Y., Hogan, T. P., Gabrieli, J. D. E., & Gaab, N. (2021). Multifactorial pathways facilitate resilience among kindergarteners at risk for dyslexia: A longitudinal behavioral and neuroimaging study. Developmental Science, 24(1), e12983. 10.1111/desc.12983

Zuk, J., Yu, X., Sanfilippo, J., Figuccio, M. J., Dunstan, J., Carruthers, C., Sideridis, G., Turesky, T. K., Gagoski, B., Grant, P. E., & Gaab, N. (2021). White matter in infancy is prospectively associated with language outcomes in kindergarten. Developmental Cognitive Neuroscience, 50, 100973. 10.1016/j.dcn.2021.100973

